# The developmental origin and the specification of the adrenal cortex in humans and cynomolgus monkeys

**DOI:** 10.1101/2022.01.19.477000

**Authors:** Keren Cheng, Yasunari Seita, Taku Moriwaki, Kiwamu Noshiro, Yuka Sakata, Young Sun Hwang, Toshihiko Torigoe, Mitinori Saitou, Hideaki Tsuchiya, Chizuru Iwatani, Masayoshi Hosaka, Toshihiro Ohkouchi, Hidemichi Watari, Takeshi Umazume, Kotaro Sasaki

## Abstract

Development of the adrenal cortex, a vital endocrine organ, originates in the adrenogonadal primordium, a common progenitor for both the adrenocortical and gonadal lineages in rodents. In contrast, we find that in humans and cynomolgus monkeys, the adrenocortical lineage originates in a temporally and spatially distinct fashion from the gonadal lineage, arising earlier and more anteriorly within the coelomic epithelium. The adrenal primordium arises from adrenogenic coelomic epithelium via an epithelial-to-mesenchymal-like transition, which then progresses into the steroidogenic fetal zone via both direct and indirect routes. Notably, we find that adrenocortical and gonadal lineages exhibit distinct HOX codes, suggesting distinct anterior-posterior regionalization. Together, our assessment of the early divergence of these lineages provides a molecular framework for understanding human adrenal and gonadal disorders.

**One Sentence Summary:** Specification of the adrenal cortex occurs in adrenogenic coelomic epithelium independent of gonadogenesis in humans and cynomolgus monkeys

## INTRODUCTION

The adrenal cortex is the major source of steroid hormones that drive a plethora of critical physiologic functions. Accordingly, developmental aberrancies in adrenal cortex formation drive various congenital and adult-onset diseases. Traditionally, rodent models have been used to assess the cellular and genetic mechanisms driving mammalian adrenal development. However, critical differences between rodents and humans exist in adrenal structure, steroidogenic function and adrenal development (*1, 2*). In fact, mutated genes implicated in human adrenal anomalies do not always result in a similar phenotype in mice, or vice versa (*1*). For example, mutations in *NR5A1*, a master transcription factor of steroidogenesis, severely affect adrenal development in mice, but rarely manifest as adrenal defects in humans (*1, 3*). Thus, the translational applicability of mechanisms driving adrenal development in murine models needs to be interpreted with caution.

Mammalian gonads (ovaries and testes) also produce steroid hormones and commonalities exist between the adrenal cortex and gonads in both their steroid synthetic pathways and developmental origin. Lineage tracing studies in mice reveal that the adrenal cortex and gonads originate from a common progenitor, the adrenogonadal primordium (*4–9*). The adrenogonadal primordium expresses GATA4 and WT1, along with NR5A1, and is thought to be established through thickening of coelomic epithelium, which is a derivative of the posterior intermediate mesoderm (*6, 7*). Thereafter, the mediodorsal portion of the adrenogonadal primordium separates and migrates dorsally to become the adrenal cortex, whereas the ventral portion becomes the gonads (*6, 7, 9*). In humans, the gonads and the adrenal cortex are first recognized as morphologically distinct structures at approximately 33 days post-conception (Carnegie stage [CS]15), at which time the adrenal cortex is recognized as a condensed blastematous structure, medial to the mesonephros, referred to as the adrenal primordium (*10–13*). However, owing to the lack of genetic tracing tools, or appropriate markers to differentiate emerging gonadal and adrenocortical lineages, how and when these lineages are specified and segregated remains poorly understood in humans.

In both mice and humans, after formation of the adrenal primordium, the adrenocortical lineage forms morphologically and functionally discrete zones: the inner fetal zone and outer definitive zone (*5, 14*). In humans and non-human primates, androgens produced by the fetal zone promote the formation of the feto-placental unit, thereby playing a pivotal role in fetal development and pregnancy maintenance (*15*). The fetal zone regress postnatally, whereas the definitive zone eventually gives rise to the adult adrenal cortex (*15*). Lineage tracing studies in mice have suggested that the fetal zone and definitive zone originate from common progenitors at the early stage of adrenal primordium (*5*). However, the molecular events accompanying the bifurcation of the fetal zone and definitive zone from the earlier progenitor have not been characterized in humans.

Through high resolution lineage trajectory mapping, we now find that adrenocortical specification in primates occurs in a spatially and temporarily distinct manner from gonadogenesis thus providing essential insight for understanding human adrenogenesis and gonadogenesis.

## RESULTS

### Specification of the gonads in the coelomic epithelium

We previously demonstrated that in both mice and cynomolgus monkeys, the posterior intermediate mesoderm gives rise to KRT19^+^ coelomic epithelium, which, in turn, sequentially expresses GATA4 and NR5A1 to acquire the gonadal fate at E10.0 in mice and E31 in cynomolgus monkeys (*8*). To precisely determine the specification timing of human gonads, we analyzed serial paraffin sections of human embryos at various stages (CS12, n=1; CS13, n=3; CS14, n=1; CS15, n=3; CS16, n=2; CS17, n=1, table S1). Similar to cynomolgus monkeys, a CS12 human embryo showed WT1^+^ posterior intermediate mesoderm extending ventrolaterally and forming KRT19^+^ early coelomic epithelium, which was demarcated from the FOXF1^+^ lateral plate mesoderm/splanchnic mesoderm (fig. S1A-D). Regional differences in posterior intermediate mesoderm maturation along the anterior-posterior axis were observed; more anterior sections exhibited increased lateral extension of early coelomic epithelium suggestive of advanced maturation, whereas the posterior-most sections showed more immature morphology within the circumscribed mass of posterior intermediate mesoderm and no extensions of early coelomic epithelium (fig. S1D). These findings are consistent with previous studies on mice and cynomolgus monkey embryos and suggest that coelomic epithelium originates from posterior intermediate mesoderm (*8*).

We next sought to determine the onset of human gonadogenesis using GATA4 and NR5A1 as markers for the emerging gonads (*8, 16*). GATA4 expression was not detected in CS12-13 embryos (n=4) (fig. S1D-F). A weak GATA4 signal was first observed in a CS14 embryo within the WT1^+^NR5A1^-^ pseudostratified coelomic epithelium at the mid-posterior regions (fig. S1F, S2A, B), which we termed gonadogenic coelomic epithelium. Early in week 5 post-fertilization (wpf) (CS15, n=3), WT1^+^GATA4^+^NR5A1^+^*LHX9*^+^ nascent gonad progenitors were observed at the ventrolateral aspect of the mesonephros as a pseudostratified region within KRT18/KRT19^+^ coelomic epithelium (fig. S2A-D). NR5A1^+^ gonad progenitors were encompassed by the region expressing GATA4, which extended more broadly posteriorly and medially (fig. S2C, E, F). By late 5 wpf, the gonads became unequivocally recognizable by histology (CS16, n=2) (fig. S2A, B). These findings reveal that human gonadogenesis is initiated at CS14-15 through the sequential activation of GATA4 and NR5A1 within the posterior coelomic epithelium, similar to mice and cynomolgus monkeys (*8, 16*).

### Specification of the adrenal cortex in the adrenogenic coelomic epithelium in humans

During the immunofluorescence studies on emerging gonads, we incidentally found a WT1^+^GATA4^-^NR5A1^+^ region within the coelomic epithelium in 3-4 wpf embryos (CS12-14) (Fig. 1A, B). This region was seen only at the anterior portion of embryos (CS13, n=2) (fig. S3A), possessed markers of coelomic epithelium (KRT18^+^KRT19^+^CDH1^-^) (*8*), but was negative for a mesonephric marker, PAX2 (*17*), suggesting that it represents a portion of the coelomic epithelium (fig. S3B). Surprisingly, the earliest stage embryo in which these cells were observed was at 3 wpf (CS12, n=1), suggesting that the GATA4^-^NR5A1^+^cells emerge shortly after the establishment of coelomic epithelium (Fig. 1C). These coelomic epithelium cells gradually transitioned into the migration stage as they increased NR5A1 expression and lost WT1 and KRT19 expression, and eventually organized into the tightly-packed adrenal primordium (organization stage) (5wpf, CS16) (Fig. 1C, D, fig. S3C, D) (*10, 12, 18*). Accordingly, these cells express key adrenocortical markers, *NR5A1*, *STAR*, *NR0B1* and *WNT4,* but lack the gonadal markers, *GATA4* and *LHX9* (Fig. 1B). Thus, in humans, the adrenocortical lineage is first specified within a portion of the anterior coelomic epithelium, which we designate the adrenogenic coelomic epithelium. Moreover, the adrenocortical and gonadal lineages are specified independently, as gonadogenic coelomic epithelium did not appear until CS14, and was only observed in the posterior sections where GATA4^-^NR5A1^+^ adrenogenic coelomic epithelium was not detected (fig. S4A). At CS15-16 when gonad progenitors appear, the WT1^-^GATA4^-^NR5A1^bright+^ adrenal primordium had already formed an organized mass, medial to the mesonephros, and were clearly separated from the WT1^+^GATA4^+^NR5A1^weak+^ emerging gonad progenitors (fig. S4B, C). Moreover, these two lineages exhibited a distinct distribution pattern along the anterior-posterior axis; the gonads extending more posteriorly than adrenal primordium (CS15, n=3) (fig. S4B).

**Fig. 1.**
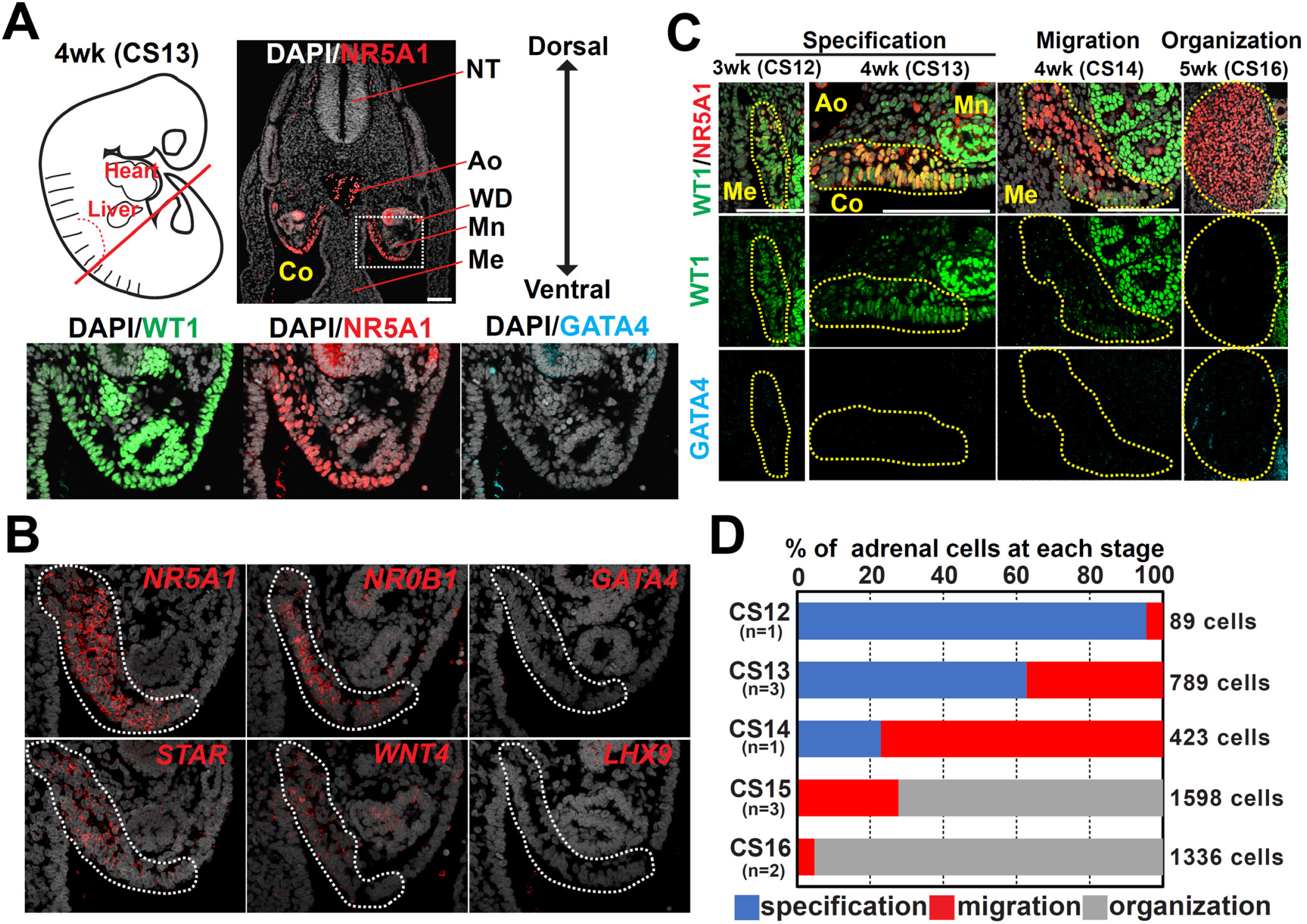
Specification of the adrenocortical lineage in humans. (A) (top left) Schematic of a human embryo at 4 wpf (CS13), with a red line indicating the approximate plane where the transverse sections were made for immunofluorescence (IF) analysis. (top right) IF image of the embryo for NR5A1 (red) merged with DAPI (white). (bottom) Magnified IF images of the region outlined by a dotted line for WT1 (green), NR5A1 (red) and GATA4 (cyan), merged with DAPI. Ao, aorta; Co, coelom; Me, mesentery; Mn, mesonephros; NT, neural tube; WD, Wolffian duct. Bars, 100 μm. (B) In situ hybridization (ISH) images of the coelomic angle (embryos: CS13) for the indicated markers (red) merged with DAPI (white). Dotted lines indicate the adrenogenic coelomic epithelium (CE). Bar, 100 μm. (C) IF images of the coelomic angle at the indicated stages for WT1 (green), NR5A1 (red) and GATA4 (cyan). Merged images for WT1, NR5A1 and DAPI (white) are shown at the top. The yellow dotted line encircles the adrenogenic CE or adrenal primordium. Bars, 100 μm. (D) Percentage of adrenocortical cells at the indicated morphogenetic stages (specification, migration and organization) for each embryonic stage (Carnegie stage [CS] 12–16). Numbers of embryos and counted cells are shown.

Previous studies in mice showed that germ cells required for eventual gamete production first migrate into the adrenogonadal primordium and redistribute into the gonads after segregation of the gonads and adrenal primordium (*7*). In humans, although primordial germ cells migrate into both the adrenal primordium and the gonads at early stages (4-5 wpf), they were selectively lost in the adrenal cortex thereafter (6wpf). Thus, species-specific mechanisms of primordial germ cell migration, redistribution and survival govern future gamete generation within gonads (fig. S4D, E).

### Specification of the adrenal cortex in cynomolgus monkeys

To evaluate whether the mode of adrenal specification is conserved among primates, we performed immunofluorescence analyses on serial sections of cynomolgus monkey embryos (CS13-16, n=11, table S1). WT1^+^GATA4^-^NR5A1^+^ adrenogenic coelomic epithelium first emerged at E28 (CS13) within the KRT18^+^KRT19^+^CDH1^-^ PAX2^-^ coelomic epithelium (Fig. 2A-D, Fig. S5A, B). Disruption of basement membranes was noted along adrenogenic coelomic epithelium, suggestive of an active epithelial-to-mesenchymal transition (EMT)-like change (fig. S5B). Upon specification, adrenogenic coelomic epithelium underwent migration while gradually losing WT1 and KRT19 expression, and formed a tightly organized WT1^-^GATA4^-^NR5A1^+^ adrenal primordium by E31 (CS16) (Fig. 2E, fig. S5B-D). When WT1^+^GATA4^+^NR5A1^+^ gonad progenitors formed at E31, the adrenocortical cells were already organized into adrenal primordium, without spatially overlapping with the gonad (fig. S5E). These results suggest that in humans and cynomolgus monkeys, both adrenocortical and the gonadal lineages originate from the coelomic epithelium, albeit in a spatially and temporally distinct manner.

**Fig 2.**
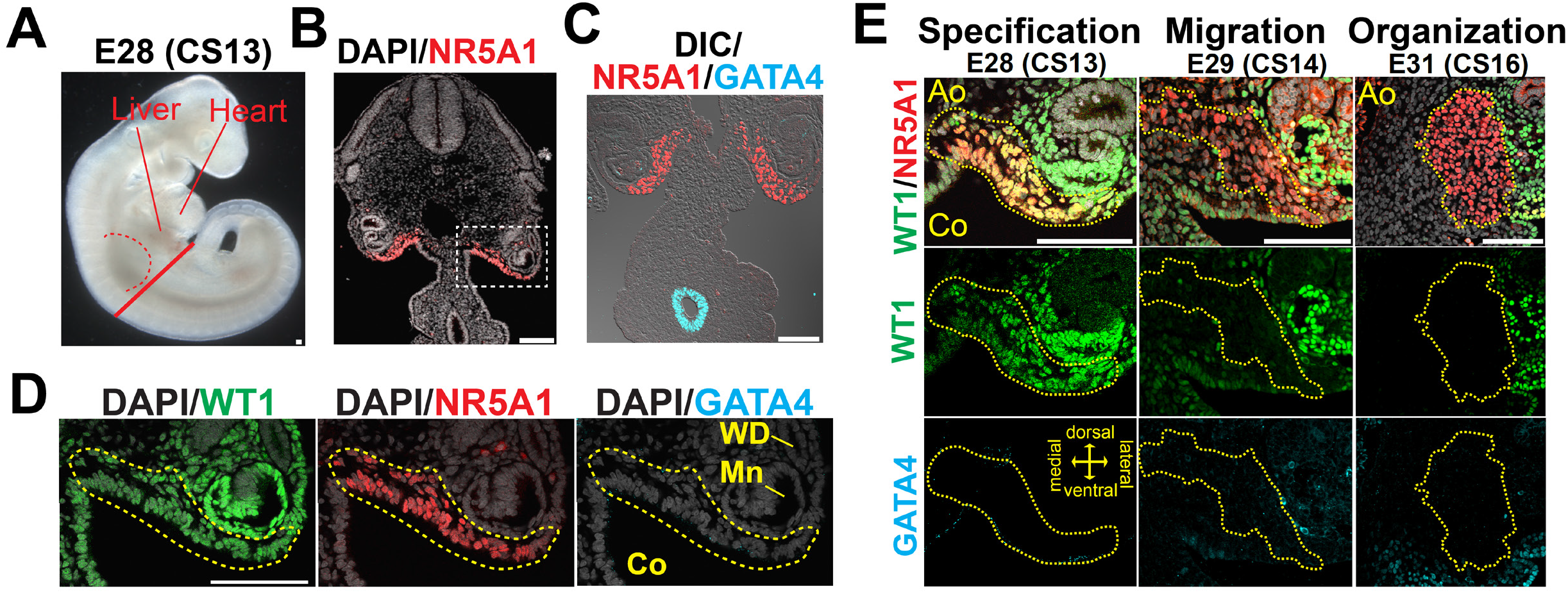
Specification of the adrenocortical lineage in cynomolgus monkeys. (A) Brightfield image of a cynomolgus (cy) embryo at E28 (CS13). The red dotted line denotes the forelimb bud. The solid red line indicates the approximate plane where the transverse sections were taken for IF analysis. (B, C) IF images of the section for NR5A1 (red) merged with DAPI (white) (B) or GATA4 (cyan), and differential interference contrast (DIC) image (white) (C). (D). Magnified images of the region outlined in (B) for WT1 (green), NR5A1 (red) and GATA4 (cyan), merged with DAPI (white). NR5A1^+^ adrenogenic CE are outlined by the yellow dotted line. Bars, 100 μm. (E) IF images of the coelomic angle of cynomolgus embryos at the indicated morphogenetic and embryonic stages for WT1 (green), NR5A1 (red) and GATA4 (cyan). Merged images for WT1, NR5A1 and DAPI (white) are shown. The yellow dotted lines outline the adrenogenic CE or adrenal primordium. Bars, 100 μm.

### Transcriptomic changes accompanying adrenogenic coelomic epithelium specification in humans

We next sought to understand the molecular events associated with the segregation and subsequent development of adrenocortical and gonadal cell lineages using single cell transcriptomics. After conducting quality control analyses (fig. S6A), we first explored adrenocortical development immediately after the specification of adrenogenic coelomic epithelium. Analyzing whole urogenital ridge at 4 wpf (CS14) by the Uniform Manifold Approximation and Projection (UMAP) revealed *WT1*^+^*KRT18*^+^*KRT19*^+^ coelomic epithelium, portions of which showed *GATA4* or *NR5A1* expression in a mutually exclusive manner (fig. S6B). Various other cell types were also annotated based on known marker genes and differentially expressed genes (fig. S6C, D, table S2).

Re-clustering of the coelomic epithelium cluster revealed two subclusters. The first cluster (colored in red) was annotated as adrenogenic coelomic epithelium given its characteristic gene expression pattern (*GATA4*^-^*LHX9*^-^*NR0B1*^+^*STAR*^+^*NR5A1*^+^) (Fig. 1A, B, 3A, B). The second cluster (colored in blue) expressed posterior *HOX* genes (e.g., *HOXA9*, *HOXD9*) and encompassed *GATA4*^+^*LHX9*^+^*NR5A1*^-^ gonadogenic coelomic epithelium, and was tentatively annotated as posterior coelomic epithelium (Fig. 3A-C and fig. S6E). Notably, the adrenogenic coelomic epithelium cluster could be further divided into two subclusters (Fig. 3D and fig. S7A). RNA velocity and pseudotime trajectory analyses revealed overall directional lineage progression from one cluster (bottom) to the other (top) (Fig. 3D). These clusters differed in expression of *NR5A1* and *WT1,* reflecting the immunophenotypic differences between NR5A1^low^WT1^high^ pre-migratory adrenogenic coelomic epithelium and NR5A1^high^WT1^low^ migratory adrenogenic coelomic epithelium (fig. S7B, C). Accordingly, genes related to cell migration were upregulated in migratory adrenogenic coelomic epithelium (fig. S7B, D, table S3). Genes enriched in GO terms such as “adrenal gland development” and “cholesterol metabolic process” were upregulated along the pseudotime, suggesting initiation of expression of the adrenal steroidogenic machinery (fig. S7D). Moreover, migratory adrenogenic coelomic epithelium showed upregulation of many genes related to “extracellular matrix” as well as key transcription factors critical for EMT (e.g., *SNAI1*, *SNAI2*, *TWIST2*, *ZEB2*) (Fig. 3E, fig. S7B, D, table S3), suggesting that the EMT-like morphogenetic changes are operative during the transition (*19*). While adrenogenic coelomic epithelium expressed some of the previously identified mouse adrenal specifier genes such as *PKNOX1* (*PREP*) and *WT1*, other key specifiers, such as *CITED2*, *PBX1* or *HOXB9* were only rarely and weakly expressed at this stage(*4, 9*), suggesting a divergence of the adrenal specification genetic pathway between mice and humans (fig. S7E).

**Fig 3.**
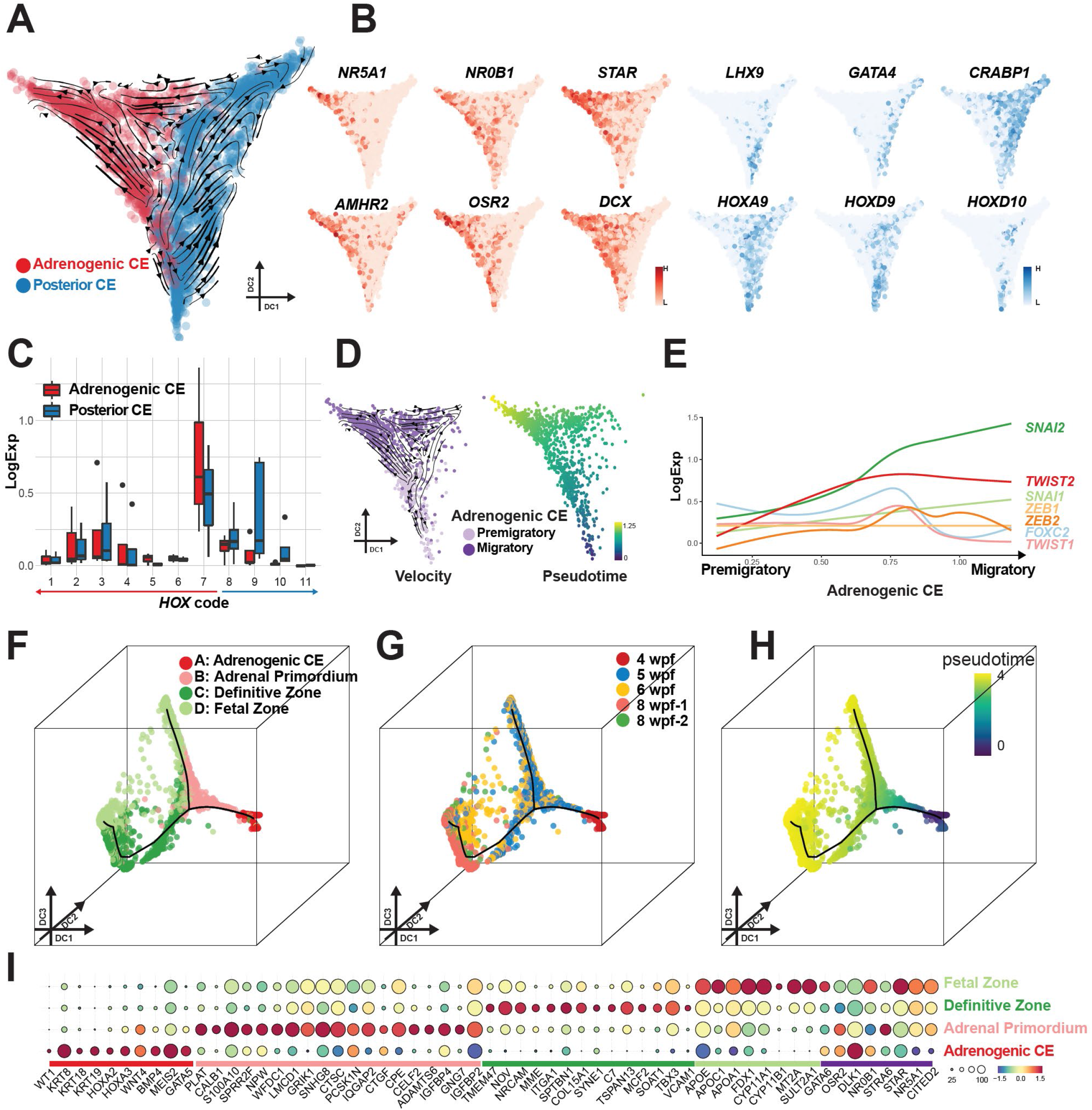
Transcriptomic dynamics accompanying adrenal specification and maturation in humans. (A) Single cell transcriptomes of the urogenital ridge at 4 wpf, projected on diffusion map with overlaid RNA velocity. Cells are colored according to cell cluster (red, adrenogenic CE; blue, posterior CE). Black lines with arrows show the velocity field. (B) Expression of key markers of adrenogenic CE (red) and posterior CE (blue). (C) Summary of *HOX* gene expression in adrenogenic CE and posterior CE. Log normalized expression of *HOX* genes belonging to the same paralogues are summarized together. Numbers 1–7 indicate anterior *HOX* genes, and 8–11 indicate posterior *HOX* genes. *HOX12* and *HOX13* are filtered out as outliers because of extremely low expression. (D) (left) Subclustering of adrenogenic CE into two cell clusters, pre-migratory and migratory adrenogenic CE, with overlaid RNA velocity. (right) Pseudotime is projected on the same plots. (E) Expression dynamics of key transcription factors associated with epithelial-to-mesenchymal transition, aligned along pseudotime as in (D). (F-H) Reclustering of the adrenocortical lineage (defined in fig. S8C) and projection of four clusters onto the diffusion map. Cells are colored according to cell cluster (F), sample origin (G) or pseudotime (H). (I) Key marker gene expression. Putative genes involved in adrenal specification and steroidogenesis are also shown at the far right, underlined by a purple bar.

### Gene expression dynamics during maturation of adrenocortical cells

We next sought to understand the dynamics of adrenocortical cell maturation after specification. UMAP plot of whole adrenals at 5-12 wpf and the urogenital ridge at 4 wpf revealed *NR5A1*^+^*STAR*^+^ fetal adrenocortical lineage (adrenogenic coelomic epithelium, cluster 3; adrenal cortex, cluster 8) (fig. S8A-C). Other lineages previously observed in fetal adrenals were also identified based on the expression of marker genes (fig. S8C, D and table S4). Re-clustering of isolated adrenocortical lineages and projection onto the diffusion map revealed a lineage trajectory in which four clusters (clusters A-D) aligned along pseudotime, which were also concordant with sample origins (Fig. 3F-H, fig. S9A, table S5). Cluster A was annotated as adrenogenic coelomic epithelium as it expressed markers of previously defined adrenogenic coelomic epithelium (e.g., *WT1*, *KRT19, NR5A1*) (Fig. 3F, I). As the lineage progressed towards cluster B, cells upregulated some steroidogenic genes (e.g., *NR5A1*, *APOA1*, *FDXR*) while they downregulated coelomic epithelium markers (e.g., *WT1*, *KRT19*, *KRT18*) (Fig. 3F-I, fig. S9B-D). Genes involved in early development and axial specification were also downregulated (fig. S9B-D). Notably, this cluster had yet to significantly upregulate genes characteristic of the fetal zone (e.g., *CYP17A1*, *SULT2A1*) or the definitive zone (e.g., *MME*, *HSD3B2*, *NOV*) (fig. S9D) and was therefore annotated as adrenal primordium. The lineage subsequently bifurcated into two branches (Fig. 3F-H). In the first branch (direct path), adrenal primordium directly fed into cluster D, which consisted predominantly of 6-8 wpf adrenals, highly expressing key enzymes required for the biosynthesis of adrenal androgens (e.g., *CYP17A1*, *SULT2A1*), features characteristic of fetal zone adrenal cells (Fig. 3F-I, fig. S9B-E, S10A-C) (*2*). The second branch (indirect path) showed transition of adrenal primordium into the fetal zone through cluster C, herein annotated as definitive zone adrenal cells based on expression of known markers of the definitive zone (e.g., *NOV*, *HSD3B2*) (Fig. 3F-I, fig. S9B-E, S10C) (*2*). Differential gene expression analysis showed a number of novel surface markers (*MME*, *ITGA1*, *NRCAM*, *TMEM47*), which might be useful for the isolation and characterization of the definitive zone (Fig. 3I, fig. S9D, S10C, table S5). We also noted that the definitive zone expressed various transcription factors/homeodomain proteins (e.g., *LEF1*, *ETV5*, *TBX3*, *HOPX*), which might play roles in maintaining the stemness of the definitive zone (Fig. 3I, S9B). Notably, fetal zone development through either the direct or indirect pathway was associated with upregulation of genes required for steroid biosynthesis while those associated with cell proliferation were downregulated, consistent with the notion that the fetal zone contains terminally differentiated steroidogenic cells (fig. S9C, E). Altogether, our findings suggest that early adrenocortical development accompanies the gradual acquisition of steroidogenic function following direct or indirect development from a common progenitor, the adrenal primordium.

Recent studies in mice suggest that a functional interaction between the adrenal capsule and the fetal adrenal cortex maintains functional zonation through signaling mediated by the Hedgehog and WNT pathways (*14*). Consistent with this notion, we found that Hedgehog ligands are expressed in the adrenal cortex while their receptors and downstream genes are expressed in the capsule. In contrast, WNT ligands (i.e., *WNT4* and *RSPO3*) were predominantly expressed in the capsule, whereas their receptors and downstream target genes are preferentially expressed in the adrenal cortex (fig. S11A-E). These findings suggest that similar interactions between the capsule and cortex play a role in maintaining the functional zonation of the human fetal adrenals.

### Adrenocortical and gonadal lineages bear distinct HOX codes

We next set out to explore the developmental trajectory of the early testicular lineages. To this end, clusters representing gonadal somatic cells were isolated from single cell transcriptomes of the whole testes at 5-12 wpf and the urogenital ridge at 4 wpf and projected onto a diffusion map (Fig. 4A, B, fig. S12A-E, S13A-C, table S6). These analyses revealed the lineage trajectory where *KRT19*^+^*WT1*^+^*GATA4*^+^*NR5A1*^-^ gonadogenic coelomic epithelium (cluster 15, predominantly 4 wpf) first progressed into *KRT19*^+^ *WT1*^+^*GATA4*^+^*NR5A1*^+^ gonad progenitors (cluster 11, predominantly 5 wpf), which subsequently bifurcate into two lineages: *SRY*^+^ Sertoli cell progenitors (cluster 8)/*AMH*^+^ Sertoli cells (cluster 3) and *ARX*^+^ interstitial progenitor (cluster 1)/*INSL3*^+^ fetal Leydig cell lineages (cluster 10) (Fig. 4A, B, fig. 12E, 13A-G).

**Fig 4.**
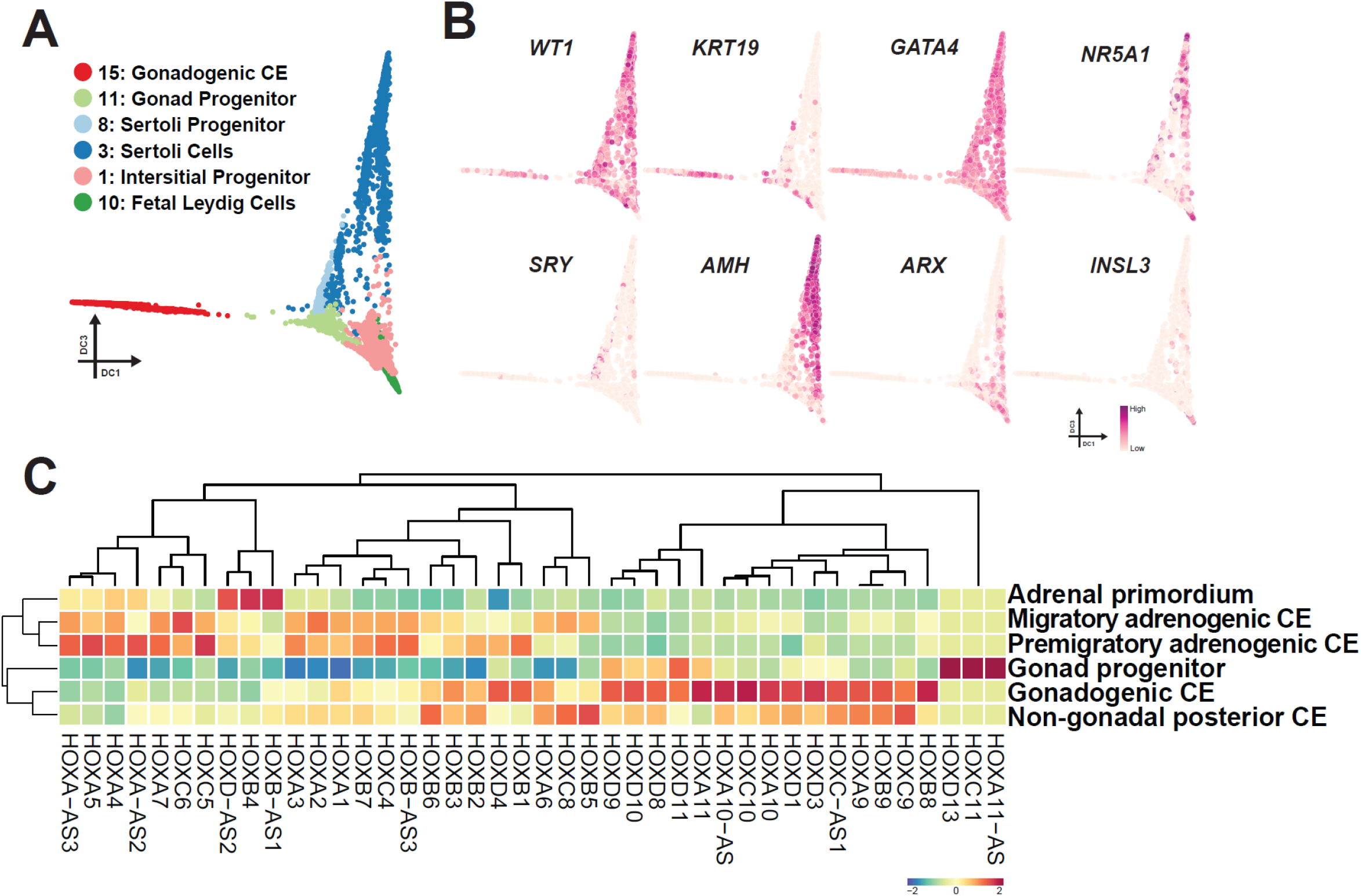
Transcriptomic dynamics during the maturation of adrenocortical cells. (A) Diffusion map showing the trajectory during human gonad development. Cells are colored according to cell cluster as in fig. S12C. (B) Key marker genes used for annotation of cell types projected on the diffusion map in (A). (C) Averaged expression values of *HOX* genes in the indicated cell clusters. *HOX* gene expression is Z-score normalized by each column, and cell clusters and genes are hierarchically clustered.

Our transcriptome analysis and annotation of cell types present in the early adrenocortical or gonadal lineages allowed us to interrogate the potential relationship between these ontologically related lineages (fig. 13H-J). Pairwise correlation analyses of cell types present in these two lineages showed an overall high degree of transcriptomic correlation (fig. 13I). Unbiased hierarchical clustering revealed gonad progenitors clustered with gonadogenic coelomic epithelium rather than adrenogenic coelomic epithelium (migratory and pre-migratory), further lending support to their distinct origins (fig. 13I). Moreover, these combined analyses reveal that adrenocortical and gonadal lineages exhibit distinct expression patterns of *HOX* genes; adrenocortical lineages expressed predominantly anterior *HOX* gene paralogues (*HOX1-7*), whereas gonadal lineages had high levels of posterior *HOX* genes (*HOX 8-13*) with only a modest expression of anterior *HOX* genes (Fig. 4C, fig. S13I-K). Altogether, these findings further support that the conclusion that adrenal and gonadal lineages have distinct origins characterized by differential expression of anterior versus posterior *HOX* codes in their early progenitors, respectively.

## DISCUSSION

In this study, we provide evidence that, in contrast to mice, the adrenal cortex in humans and cynomolgus monkeys originates in a spatially, temporally, and phenotypically distinct manner from that of the gonads (Fig. 5). Specifically, through histologic and transcriptomic analyses of early human gonadogenesis, we found that the gonad is established from the posterior coelomic epithelium at 4-5 wpf through the sequential activation of GATA4 (4 wpf, CS14) and NR5A1 (5 wpf, CS15), similar to mice and cynomolgus monkeys (Fig. 5, fig. S1) (*8*). A previous scRNA-seq study suggested that NR5A1 is not activated in the early gonad, and GATA4 only appears at 6-7 wpf when testicular cord formation is initiated (*20*). However, this discrepancy may be explained by either differences in sensitivity of detection and/or potential differences in allocation of the fetal ages relative to our study, in which the onset of gonadogenesis was more in accord with the previous immunofluorescence/histologic studies on human and monkey embryos (*8, 11, 18*). Serendipitously, during this analysis we found that the adrenocortical lineage in humans is first specified within a portion of the coelomic epithelium that is more anterior to the gonadogenic coelomic epithelium. Moreover, gonadogenic coelomic epithelium appeared later in development than did adrenogenic coelomic epithelium, and was only observed in the posterior sections where adrenogenic coelomic epithelium was not detected. Thus our findings reveal that, in contrast to mice, the adrenocortical and gonadal lineages are specified independently in humans.

**Fig 5.**
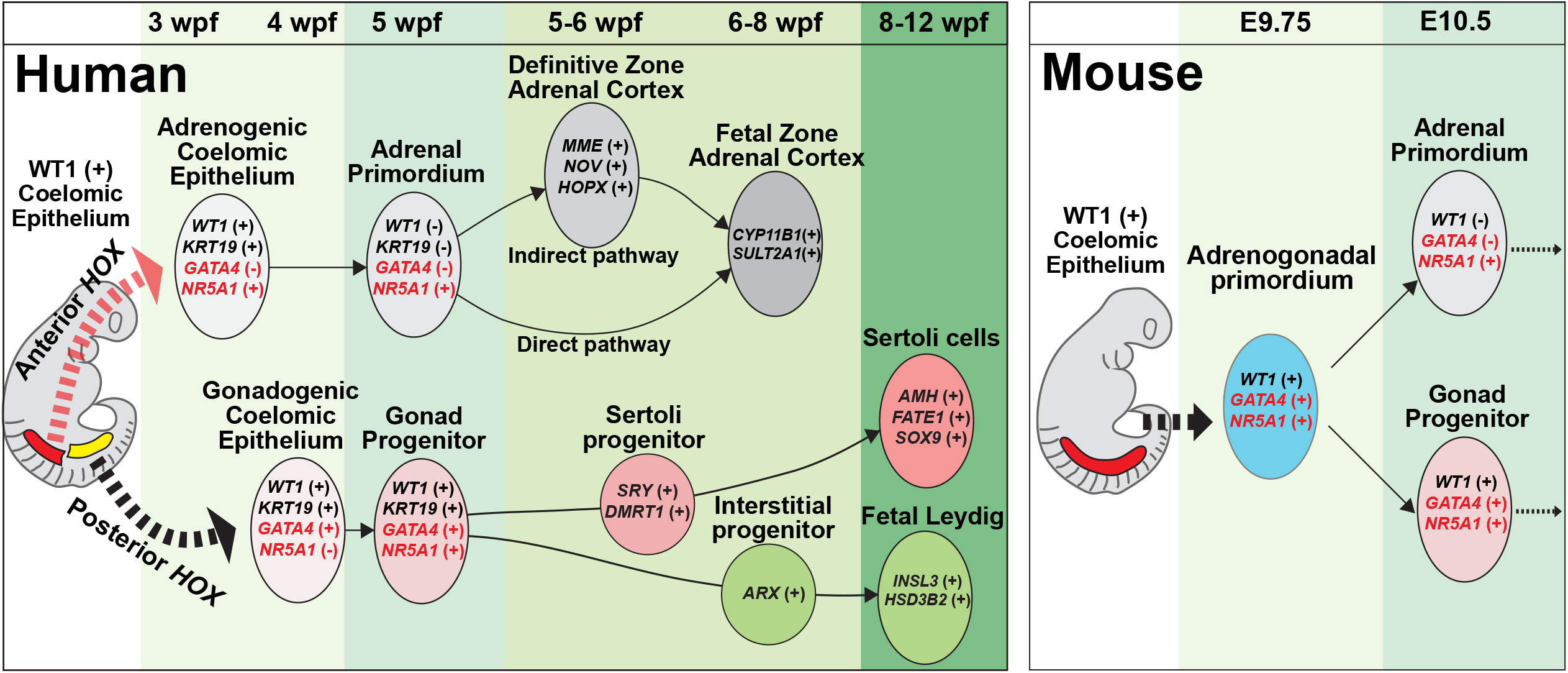
A model of adrenal and gonadal development in humans and mice. Schematic representation of a model of adrenal and gonadal development in humans (left) and mice (right).

The origin of the fetal and definitive zones in the human fetal adrenal cortex has long been a source of controversy (*2, 13, 21, 22*). Our lineage trajectory analyses of human adrenal cortex transcriptomes suggest the presence of a shared progenitor of the fetal and definitive zones (i.e., adrenal primordium), consistent with lineage tracing studies in mice (*5*). However, our trajectory analysis also revealed that the definitive zone in humans can further differentiate into the fetal zone with similar steroidogenic characteristics to those derived from the direct pathways. This finding is consistent with a previous model in which the fetal zone is derived from the definitive zone through continuous centripetal migration of definitive zone cells (migration theory) (*2*). Our unifying model of early adrenal development will therefore shed new insight into the molecular ontogeny of functional zonation in human adrenal glands (Fig. 5).

We found that the adrenocortical and gonadal lineages exhibit different expression patterns of *HOX* genes, which confer regional identity along anterior-posterior axis through a “*HOX* code” that results from timed activation of *HOX* paralogous genes from the 3’ to 5’ direction as nascent mesodermal cells emigrate from the posterior growth zone (*23, 24*). While the adrenals preferentially express anterior *HOX* genes (i.e., *HOX1- 7*), the gonads express more posterior *HOX* genes (i.e. *HOX8-13*) (Fig. 4C). Thus, by the time the adrenals and the gonads are specified within the anterior and posterior coelomic epithelium, respectively, a combination of their distinct regional identity conferred by the HOX code combined with inductive signals likely establishes the unique transcriptional regulatory modules that direct adrenal or gonad-specific cell fates. In support, the Hox/Pbx1/Prep1 complex appears to be critical for the initiation of adrenal development upon segregation from the adrenogonadal primordium in mice (*4*). Thus our findings suggest that distinct induction of human adrenal and gonadal fate in vitro might be achieved through modulating signals for anterior-posterior regionalization (e.g. canonical WNT signaling) as exemplified by the recent success in the induction of anteriorly- derived ureteric bud and posteriorly-derived metanephric mesenchyme (*25, 26*).

In summary, our findings reveal the distinct origin of the human and infra- human primate adrenal cortex and gonads and provide an example of the divergence of organ morphogenesis between species. Moreover, the molecular details of early lineage diversification of human and cynomolgus monkey adrenals and gonads will serve as both a framework for understanding the molecular pathology of human disorders (e.g., disorders of sex development, primary adrenal insufficiency), and also provide essential insight into the future reconstitution of these lineages in vitro from pluripotent stem cells to advance disease modeling and regenerative medicine.

## Materials and Methods

### Collection of human embryo samples

Urogenital organs at 3–8 wpf were obtained from donors who had provided informed consent and underwent elective abortion at the University of Pennsylvania, Ohkouchi Obstetrics and Gynecology Clinic and Fukuzumi Obstetrics and Gynecology Clinic. Embryo ages were determined through ultrasonographic measurement of the crown lump length. All experimental procedures were approved by the IRBs at the University of Pennsylvania (#832470) and Hokkaido University (19–066). In most embryos, the sex was determined with sex-specific PCR performed on genomic DNA isolated from trunk tissues with primers specific to the ZFX/ZFY loci (*8, 27*). Embryos were dissected in RPMI-1640 (Roche). Human embryos used in this study are listed in Table S1.

### Collection of cynomolgus monkey embryo samples

All procedures in cynomolgus monkeys were approved by the Animal Care and Use Committee of the Shiga University of Medical Science. The assisted reproductive technologies in cynomolgus monkeys, including oocyte collection, intra-cytoplasmic sperm injection, pre-implantation embryo culture and transfer of pre-implantation embryos into recipient mothers, were as previously reported (*28*). The light cycle comprised 12 h of artificial light from 8 a.m. to 8 p.m. Water was made available ad libitum. The temperature and humidity in the animal rooms were maintained at 23–27 °C and 45–55%, respectively. Implanted embryos were scanned with transabdominal ultrasound and recovered by cesarean section under full anesthesia. A total of ten embryos were harvested, ranging in embryonic age from E28 to E32 (CS13–16 embryos; n=6 male, 3 female and 1 unknown). We included samples obtained in previous studies in the analysis (*8, 27*). The recipient females were maintained after surgery. The sex of each sampled fetus was determined by sex-specific PCR with primers targeting the ZFX/ZFY loci (*8, 27*). Cynomolgus monkey embryos used in this study are listed in Table S1.

### Antibodies

The primary antibodies used in this study included goat anti-FOXF1 (R&D Systems, AF4798; RRID:AB_2105588), goat anti-GATA4 (Santa Cruz Biotechnology, sc-1237, RRID:AB_2108747), mouse anti-GATA4 (Santa Cruz Biotechnology, sc-25310; RRID:AB_2108747), mouse anti-MME (Biocare Medical, CM129; RRID:AB_10579027), mouse anti-NR5A1 (Novus Biologicals, N1665; RRID:AB_1962633), mouse anti-TFAP2C (Santa Cruz Biotechnology, sc-12762; RRID:AB_667770), rabbit anti-CDH1 (Cell Signaling, 3195S; RRID:AB_2291471), rabbit anti-laminin (Abcam, ab11575; RRID:AB_298179), rabbit anti-WT1 (Abcam, ab89901; RRID:AB_2043201), rabbit anti-KRT18 (Abcam, ab133263; RRID:AB_11155892), rabbit anti-KRT19 (Abcam, ab52625; RRID:AB_2281020) and rabbit anti-PAX2 (BioLegend, 901002; RRID:AB_2734656). The secondary antibodies included Alexa Fluor 488 conjugated donkey anti-rabbit IgG (Life Technologies, A21206; RRID:AB_2535792), Alexa Fluor 488 conjugated donkey anti-mouse IgG (Life Technologies, A32766; RRID:AB_2762823), Alexa Fluor 568 conjugated donkey anti-mouse IgG (Life Technologies, A10037; RRID:AB_2534013), Alexa Fluor 568 conjugated donkey anti-rabbit IgG (Life Technologies, A10042; RRID:AB_2534017), Alexa Fluor 647 conjugated donkey anti-goat IgG (Life Technologies, A21447; RRID:AB_2535864) and Alexa Fluor 647 conjugated donkey anti-rabbit IgG (Life Technologies, A31573; RRID:AB_2536183).

### IF analyses on paraffin sections

IF analyses for human and cynomolgus monkey embryos were performed on paraffin sections. Anterior tissues (the head and the upper chest above the level of the heart) were removed before fixation to increase the perfusion with 10% neutral buffered formalin. Samples were kept in ice cold RPMI medium during dissection and placed in formalin fixative within 1 hr (cynomolgus embryos) or 2 hrs (human embryos) after surgical recovery of the embryos. Samples submerged in formalin were incubated ∼24 hrs at room temperature with gentle rocking. After dehydration, tissues were embedded in paraffin, serially sectioned at 4 mm thickness with a microtome and placed on glass slides (Platinum Pro). Embryos were oriented perpendicularly to the surface of the mold to obtain transverse sections. For one embryo at 8 wpf, the adrenal gland was isolated and fixed in 10% neutral formalin before being embedded in paraffin, oriented perpendicularly to the surface of the mold. The embryos embedded in paraffin blocks were serially sectioned at 4 μm thickness with a microtome (Leica RM2035) and placed on microscopic glass slides (*Superfrost Plus*, Fisher Scientific). Paraffin sections were then de-paraffinized with xylene. Antigens were retrieved by treatment of sections with HistoVT one (Nacalai Tesque) for 35 min at 90°C and then for 10 min at room temperature. The slides were washed twice with PBS, then incubated with blocking solution (5% donkey serum, 0.2% Tween 20 and 1× PBS) for 1 hr at room temperature. The primary antibody incubation was performed overnight at 4°C, and slides were washed with PBS six times (20 min each), then incubated with secondary antibodies in bocking solution and 1 μg/mL DAPI for 50 min. Slides were subsequently washed six times with PBS before being mounted in Vectashield mounting medium (Vector Laboratory) for confocal microscopy analysis (Leica, SP5-FLIM inverted). Confocal images were processed in Leica LasX (version 3.7.2).

### ISH on paraffin sections

ISH on formalin-fixed paraffin-embedded sections was performed with a ViewRNA ISH Tissue Assay Kit (Thermo Fisher Scientific) with gene-specific probe sets for human *NR5A1*, *LHX9*, *GATA4*, *WNT4*, *STAR*, *NR0B1*, *NOV*, *RSPO3* and *PDGFRA*. Experiments were conducted according to the manufacturer’s instructions (incubation with pretreatment buffer for 12 min, protease treatment for 6 min 30 sec and use of FastRed as a chromogen). Slides were counterstained with 1 μg/mL DAPI for 50 min before being mounted in Vectashield mounting medium for confocal microscopic analysis.

### Staging of embryos and histologic quantification of adrenals, gonads and germ cells

Embryos were staged according to the Carnegie stage whenever possible (*29*). All image analyses were performed in ImageJ. For quantification of adrenocortical cells, sections containing NR5A1^+^GATA4^-^ adrenogenic CE/adrenal primordium were first determined with IF conducted on every five sections (∼20 μm interval). Then the regions containing adrenogenic CE/adrenal primordium were partitioned into quadrants, and at least one section per quadrant was analyzed to determine the number of cells. Cells in specification stage were defined as NR5A1^+^GATA4^-^ cells within the CE consisting of a three cell layer thickness. Cells at the migration stage were defined as NR5A1^+^GATA4^-^ cells not in the CE and not forming a cluster of a size meeting the criteria for the organization stage. Cells at the organization stage were defined as NR5A1^+^GATA4^-^ cells forming >20 coherent cell clusters (<4 μm distance between cells).

### 10x Genomics single-cell RNA-seq library preparation

Fetal urogenital organs at 4–8 wpf (four adrenals [5 wpf, n=1; 6 wpf, n=1; 8 wpf, n=2], five testes [5 wpf, n=2; 6 wpf, n=1; 8 wpf, n=2; 12 wpf, n=1] and one whole urogenital ridge [4 wpf, n=1]) were used for scRNA-seq with a Chromium Single Cell 3’ Reagent Kit (v3 chemistry). For preparation of 4 wpf embryos, the lateral abdominal wall and limb bud were first trimmed with forceps and ophthalmic scissors, and then the entire urogenital ridge containing the CE, mesonephros and a portion of the proximal mesentery was isolated for downstream processing. For the 5–8 wpf samples, under a dissection microscope, the entire adrenal glands and/or testes were dissected by removal of the surrounding fibroadipose tissues. The 5 wpf samples were small and partly disrupted during the clinical procedure; therefore, complete separation of the gonads or adrenals from the surrounding tissues could not be performed with certainty. Thus, a small portion of the surrounding tissues remained attached when samples were processed for downstream assays.

Isolated embryonic fragments were washed twice with PBS and then minced with scissors in 500 μl of 0.1% trypsin/EDTA solution, then incubated for 9 min at 37°C with gentle pipetting every 3 min. After quenching of the reaction by addition of 500 μl of STO medium, cell suspensions were strained through a 70 µm nylon cell strainer and centrifuged for 220 g for 5 min. Cell pellets were resuspended in 0.1% BSA in PBS and counted. All samples were stained with trypan blue and confirmed to be >80% viable. Cells were loaded into Chromium microfluidic chips and used to generate single cell gelbead emulsions with a Chromium controller (10x Genomics) according to the manufacturer’s protocol. Gelbead emulsion-RT was performed with a T100 Touch Thermal Cycler (Bio-Rad). All subsequent cDNA amplification and library construction steps were performed according to the manufacturer’s protocol. Libraries were sequenced with a NextSeq 500/500 high output kit v2 (150 cycles) (FC-404-2002) on an Illumina NextSeq 550 sequencer.

### Mapping reads of 10x Chromium scRNA-seq and data analysis

Raw data were demultiplexed with the mkfastq command in Cell Ranger (v2.1.0) to generate Fastq files. Trimmed sequence files were mapped to the reference genome for humans (GRCh38) provided by 10x Genomics. Read counts were obtained from outputs from Cell Ranger.

Secondary data analyses were performed in Python with Scanpy 1.8.1 or in R (v.3.6.1) with the Seurat (v.4.0), ggplot2 (v.3.3.2), gplots (v.3.0.3), qvalue (v.2.18.0), maptools (v.0.9-9), genefilter (v.1.68.0), rgl (v.0.100.54), dplyr (v.0.8.3) and matrix (v.1.2-18) packages and Excel (Microsoft). UMI count tables were first loaded into R by using the Read10x function, and Seurat objects were built from each sample. Cells with fewer than 200 genes, an aberrantly high gene count above 7000 or a percentage of total mitochondrial genes >15% were filtered out. Of the ∼89477 cells for which transcriptomes were available, 72257 cells passed quality-control dataset filters and were used in downstream analysis. We detected ∼3616 median genes/cell at a mean sequencing depth of ∼60509 reads/cell (fig. S6A). Samples were combined, and the effects of mitochondrial genes, cell cycle genes and batches, were regressed out with SCTransform during normalization in Seurat and then converted to log2 (CP10M+1) values. Mitochondrial genes and cell cycle genes were excluded during cell clustering, dimensional reduction and trajectory analysis. Cells were clustered according to a shared nearest neighbor modularity optimization based clustering algorithm in Seurat. Clusters were annotated on the basis of previously characterized marker gene expression with the FeaturePlot function and the gene expression matrix file, and cluster annotation was generated for downstream analyses. Dimensional reduction was performed with the top 3000 highly variable genes and the first 30 principal components with Seurat.

Differentially expressed genes (DEGs) in different clusters were calculated with Seurat findallmarkers, with thresholds of an average log2 FC of approximately 0.25 and a *p*-value <0.01. Developmental trajectories of cells were simulated with the first 30 PCs and 30 diffusion components by DiffusionMap in destiny and scanpy. Trajectory principal lines were fitted with ElPiGraph.R. Pseudotime was calculated with dpt in destiny. RNA velocity matrices were generated with velocyto 0.17, then analyzed with scVelo 0.25. DEGs between two groups in scatterplots were identified with edgeR 3.34.1 through application of a quasi-likelihood approach and the fraction of detected genes per cell as covariates. The DEGs were defined as those with FDR <0.01, a *p*-value <0.01 and a log2- fold change >1. The cell cycle was analyzed with CellCycleScoring in Seurat. Data were visualized with ggplot2 and pheatmap. Genes in the heatmap were hierarchically clustered according to Euclidean distance, scaled by row and then visualized with pheatmap. HOX code scoring was calculated with area under the curve methods in AUCell 1.14.0. Gene ontology enrichment was analyzed with DAVID v6.8.

## ACKNOWLEDGMENTS

We thank L. King, K. Zaret and J. Strauss III for carefully reviewing the manuscript and providing insightful comments. We thank Women’s Heath and Clinical Research Center and Tumor Tissue and Biospecimen Bank at the University of Pennsylvania for human sample collection and Comparative Pathology Core at the University of Pennsylvania School of Veterinary Medicine for making histologic sections. We acknowledge K. Susztak, and K. Murata for allowing us to use 10X Genomics Chromium, J. Wang for kindly sharing their microtome with us and I. Okamoto for technical assistance for immunofluorescence studies. We thank members of Sasaki lab for the discussion of this study.

## Authors contributions

K.S. conceived the project and designed the overall experiments. K.S. and K.C. wrote the manuscript. K.S., K.C., T.M. Y.Sakata conducted the overall experiments and analyzed the data. Y.Seita., K.C., Y.S.H., T.T. contributed to the processing and analyses of scRNA-seq data. Y.Seita, K.N., M.H., T.O., H.W., T.U. contributed to the processing of human embryo materials. K.S., M.S., H.T., C.I. contributed to the isolation of cynomolgus monkey embryos.

## Data and materials availability

All data are available in the manuscript or the supplementary material.

## SUPPLEMENTARY MATERIALS

**Figs. S1** to S13 **Tables S1** to S6

## Supplementary Materials

### Supplementary Figure Legends

**Fig. S1.**
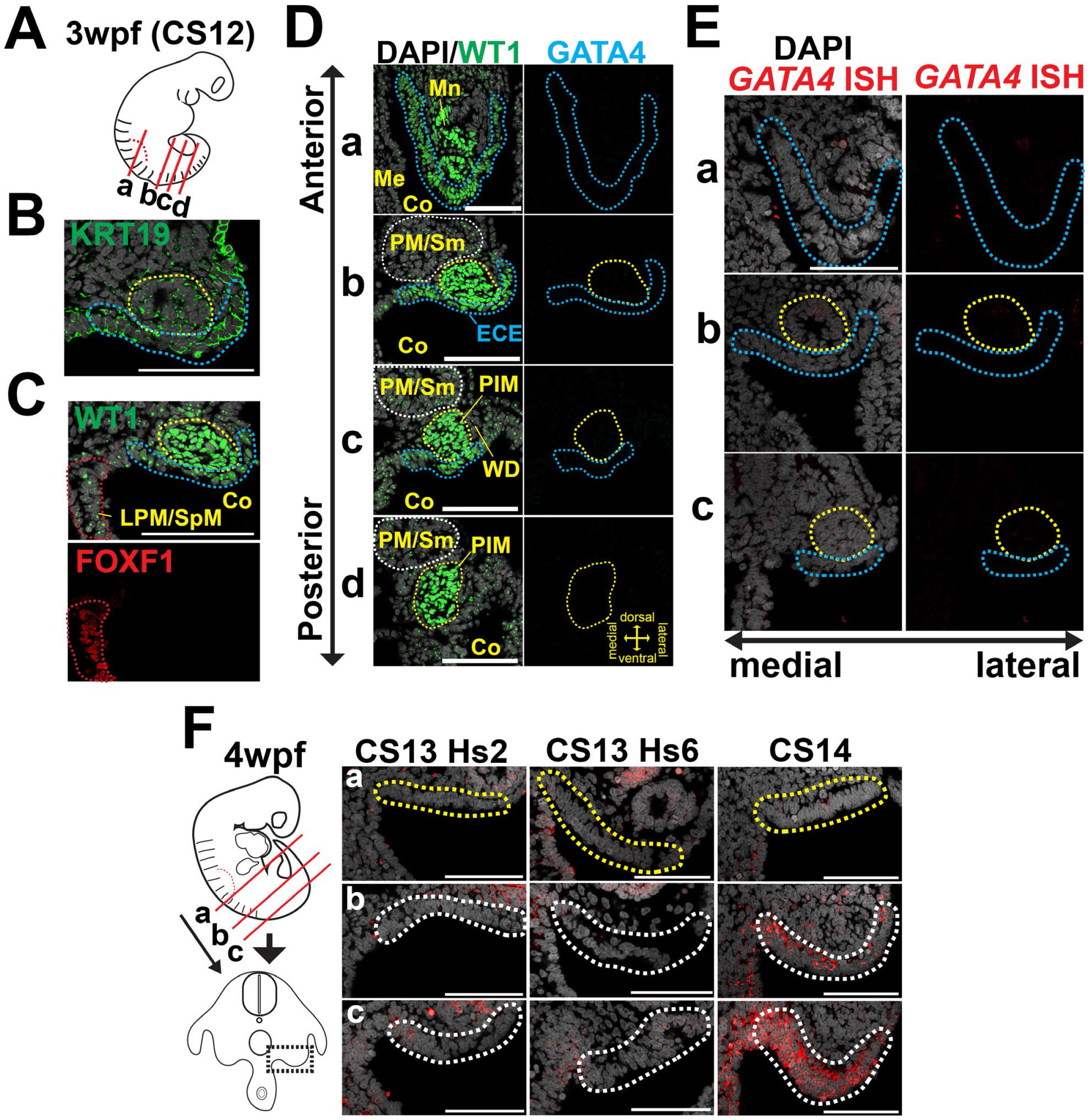
Specification of coelomic epithelium in humans. (A) Schematic of a human embryo at 3 wpf (CS12), with red lines indicating the approximate planes where the sections were taken in (B–E). (B) IF image of a transverse section taken at level b of the embryo in (A) for KRT19 (green) merged with DAPI (white). Posterior intermediate mesoderm and early coelomic epithelium (CE) are indicated by yellow and blue dotted lines, respectively. Left, medial; right, lateral. Bar, 100 μm. (C) IF images of the neighborhood section of (B) for WT1 (green) merged with DAPI (white) (top) or FOXF1 (red) (bottom). FOXF1^+^ LPM/SpM (lateral plate mesoderm/splanchnic mesoderm) is outlined by a red dotted line. Co, coelom. Bar, 100 μm. (D) IF images of the sections taken at the indicated levels, stained with WT1 (green), GATA4 (cyan) and DAPI (white). The white dotted lines outline paraxial mesoderm/somites (PM/Sm). Ms, mesonephros; Me, mesentery; WD, Wolffian duct. Bar, 100 μm. (E) ISH images of sections taken at the indicated levels shown in (A) for *GATA4* (red) merged with DAPI (white). (F) (left) Schematic of a 4 wpf human embryo and a transverse section. Approximate axial levels where transverse sections were taken for ISH are indicated by red lines. (right) ISH images of embryos at the indicated stages for *GATA4* (red) merged with DAPI (white). The region highlighted by the black dotted line in the left schematic are shown. NR5A1^+^ or NR5A1^-^ CE confirmed by IF on adjacent sections are outlined by yellow or white dotted lines, respectively. Of note, *GATA4*^+^ regions are present within CE overlying the ventral aspect of the mesonephros and proximal mesentery. Bar, 100 μm.

**Fig. S2.**
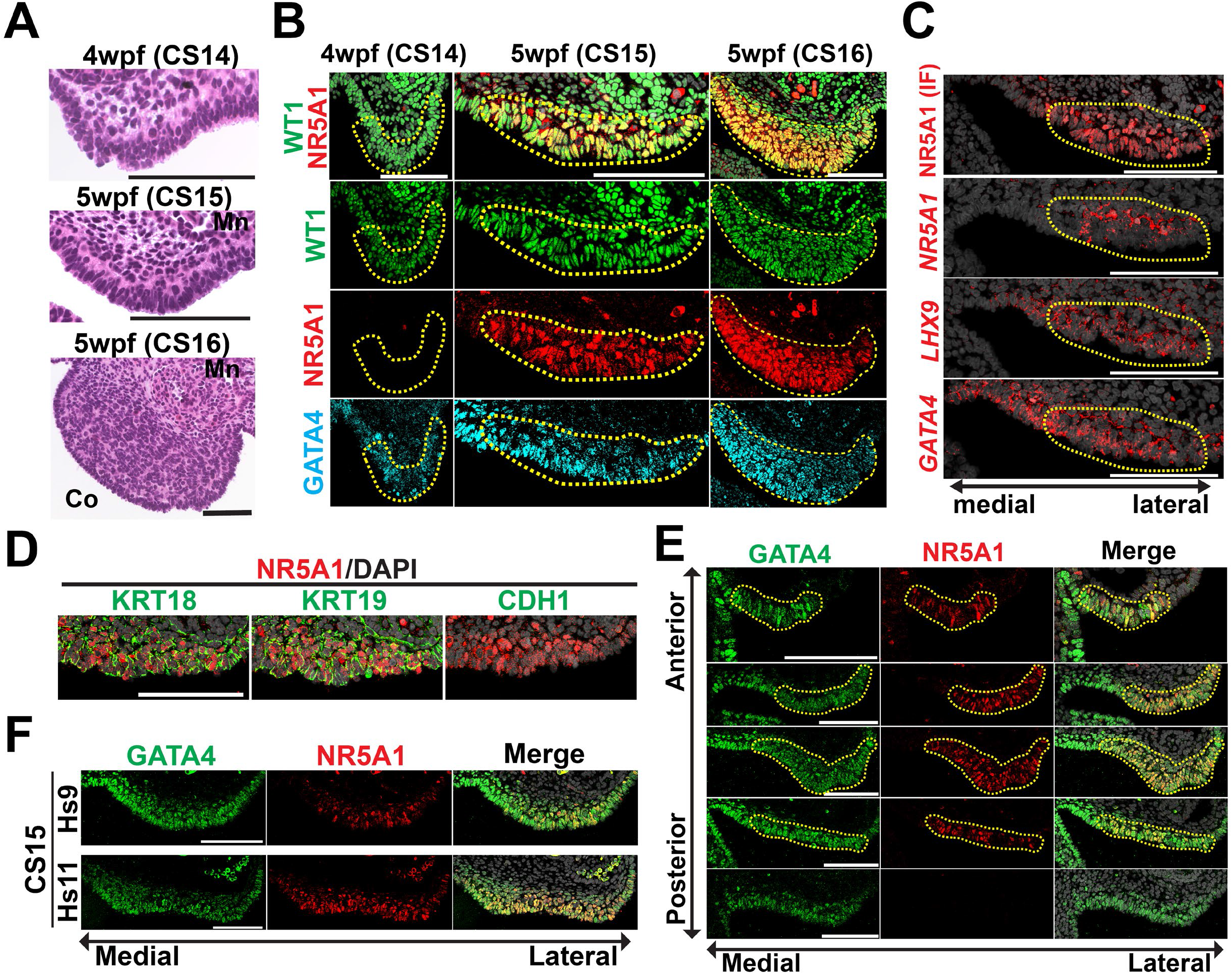
Specification of the gonads in humans. (A) Representative H&E images of developing human gonads at the indicated stages. Of note, pseudostratification of CE is already evident at CS14. Left, medial; right, lateral. Bar, 100 μm. (B) IF images of emerging gonads (outlined by yellow dotted lines) from posterior sections at the indicated stages for WT1 (green), NR5A1 (red) and GATA4 (cyan). Merged images of WT1 and NR5A1 with DAPI (white) are shown at the top. Left, medial; right, lateral. Bar, 100 μm. (C) ISH images of serial sections of the coelomic angle from a 5 wpf embryo (CS15) for the indicated markers (red) merged with DAPI (white). IF for NR5A1 on the neighboring section is also shown at the top. The NR5A1^+^ gonad progenitor is outlined by yellow dotted lines. Bar, 100 μm. (D) IF images of the gonad progenitor from the same embryo as in (C) for the indicated markers (green) merged with NR5A1 (red) and DAPI (white). Bar, 100 μm. (E) Transverse sections of the gonad progenitor from an embryo at 5 wpf (CS15) from different axial levels along the anterior-posterior axis, stained with GATA4 (green) and NR5A1 (red), and merged with DAPI (white). Of note, the NR5A1^+^ regions outlined by yellow lines are encompassed by GATA4^+^ regions. Bar, 100 μm. (F) IF images of the gonad progenitors from two embryos at CS15 (Hs9 and Hs11) stained with GATA4 (green) and NR5A1 (red), and merged with DAPI (white). Bar, 100 μm.

**Fig. S3.**
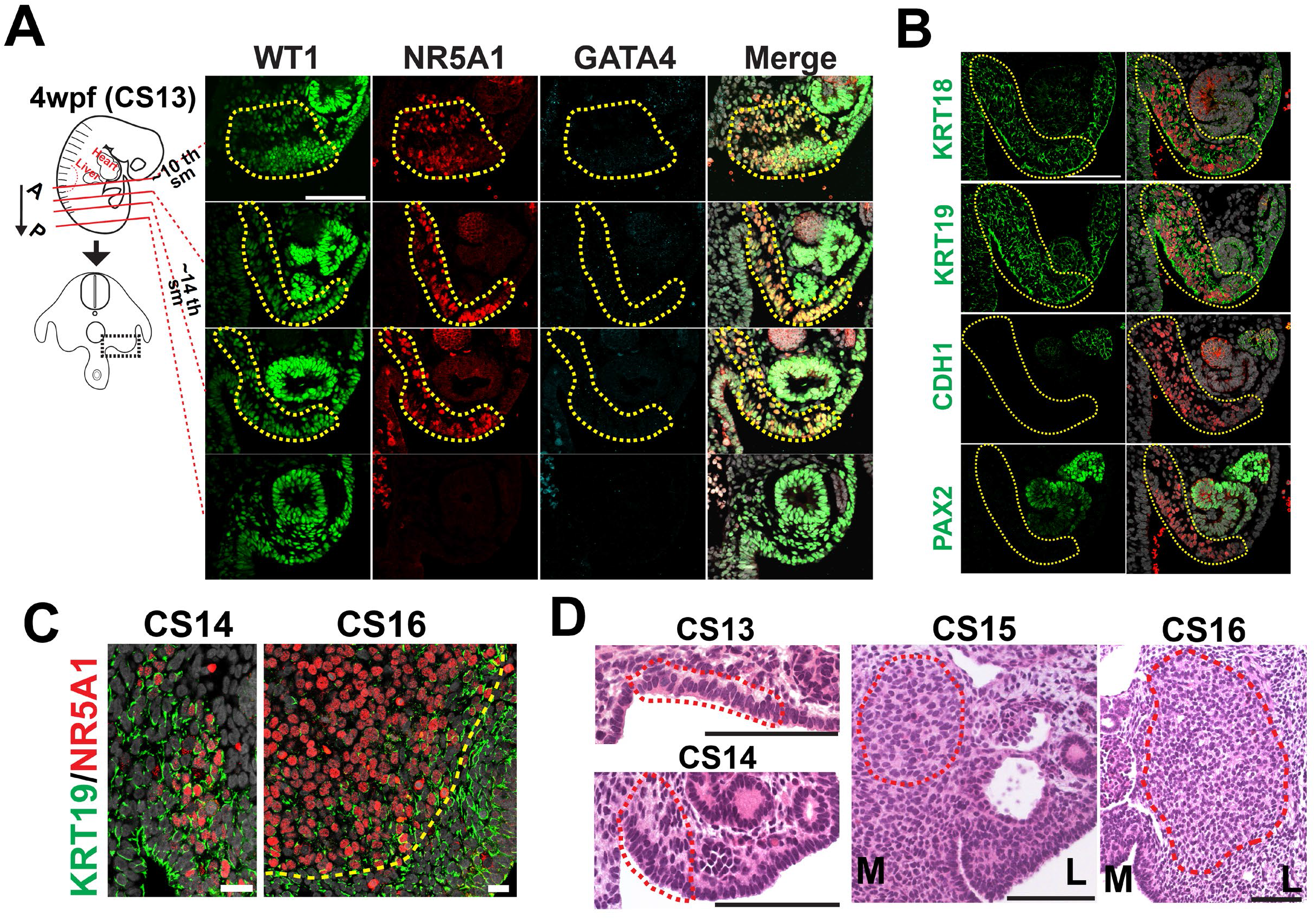
Specification of the adrenal cortex in humans. (A) (left) Schematic of a human embryo at 4 wpf (CS13), indicating planes where the transverse sections were taken. (right) IF images of these sections for WT1 (green), NR5A1 (red) and GATA4 (cyan), merged with DAPI (white). The yellow dotted lines outline the NR5A1^+^ adrenogenic CE. Of note, NR5A1^+^ adrenogenic CE is seen only at the anterior portion of the embryo (axial levels between approximately the 10^th^ somites just beneath the posterior edge of the liver/anterior limb bud and the 14^th^ somites). Bar, 100 μm. (B) IF images of the coelomic angle (embryos: CS13) for the indicated markers (green) (left). Merged images with NR5A1 (red) and DAPI (white) are shown (right). Bar, 100 μm. (C) IF images of the adrenocortical cells at the migration (CS14, left) or organization stage (CS16, yellow dotted line, right) for KRT19 (green) and NR5A1 (red), merged with DAPI (white). Bar, 20 μm. (D) H&E-stained sections showing the adrenogenic CE of predominantly pre-migratory (left top) and migratory stages (left bottom), and the adrenal primordium (CS15, middle; CS16, right), which are outlined by red dotted lines. M, medial; L, lateral. Bar, 100 μm.

**Fig. S4.**
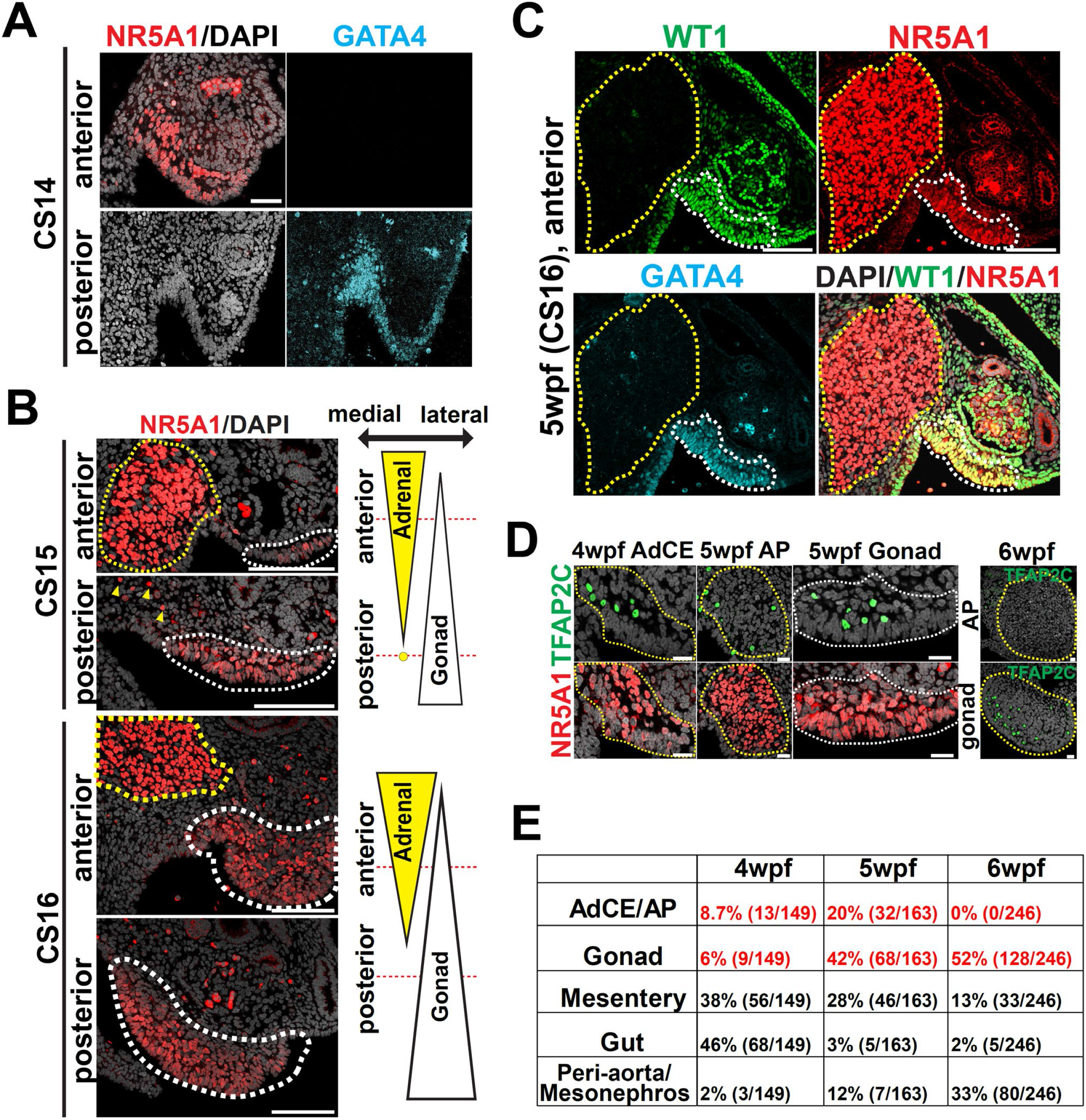
Spatial orientation of adrenocortical and gonadal lineages in humans. (A) IF images of the coelomic angle from a 4 wpf human embryo (CS14) for NR5A1 (red) and GATA4 (cyan). Merged images of NR5A1 and DAPI (white) are shown on the left. Left, medial; right, lateral. Bar, 50 μm. (B) IF images of the transverse sections taken from the indicated positions from embryos at CS15 (top) or CS16 (bottom). Merged images of NR5A1 (red) and DAPI (white) are shown. The yellow and white dotted lines outline the adrenal primordium and gonad progenitor, respectively. The yellow arrowheads indicate several scattered cells of adrenal origin. Bar, 100 μm. (C) IF images of the anterior section taken from a 5 wpf human embryo (CS16), stained with WT1 (green), NR5A1 (red), GATA4 (cyan). A merged image of WT1, NR5A1 and DAPI (white) is also shown. Bar, 100 μm. (D) IF images of human embryos (4–6 wpf, transverse sections) at the indicated regions for TFAP2C (primordial germ cells [PGC] marker, green) and NR5A1 (red), merged with DAPI (white). AdCE, adrenogenic CE; AP, adrenal primordium. (E) Distribution of PGCs in different anatomic locations of human embryos at 4–6 wpf, assessed in transverse sections stained with TFAP2C (PGC marker), NR5A1 (markers for AdCE, AP, gonads) and GATA4 (gonad marker), merged with DAPI. The percentage (number) of PGCs in each anatomic location for all counted PGCs is shown.

**Fig. S5.**
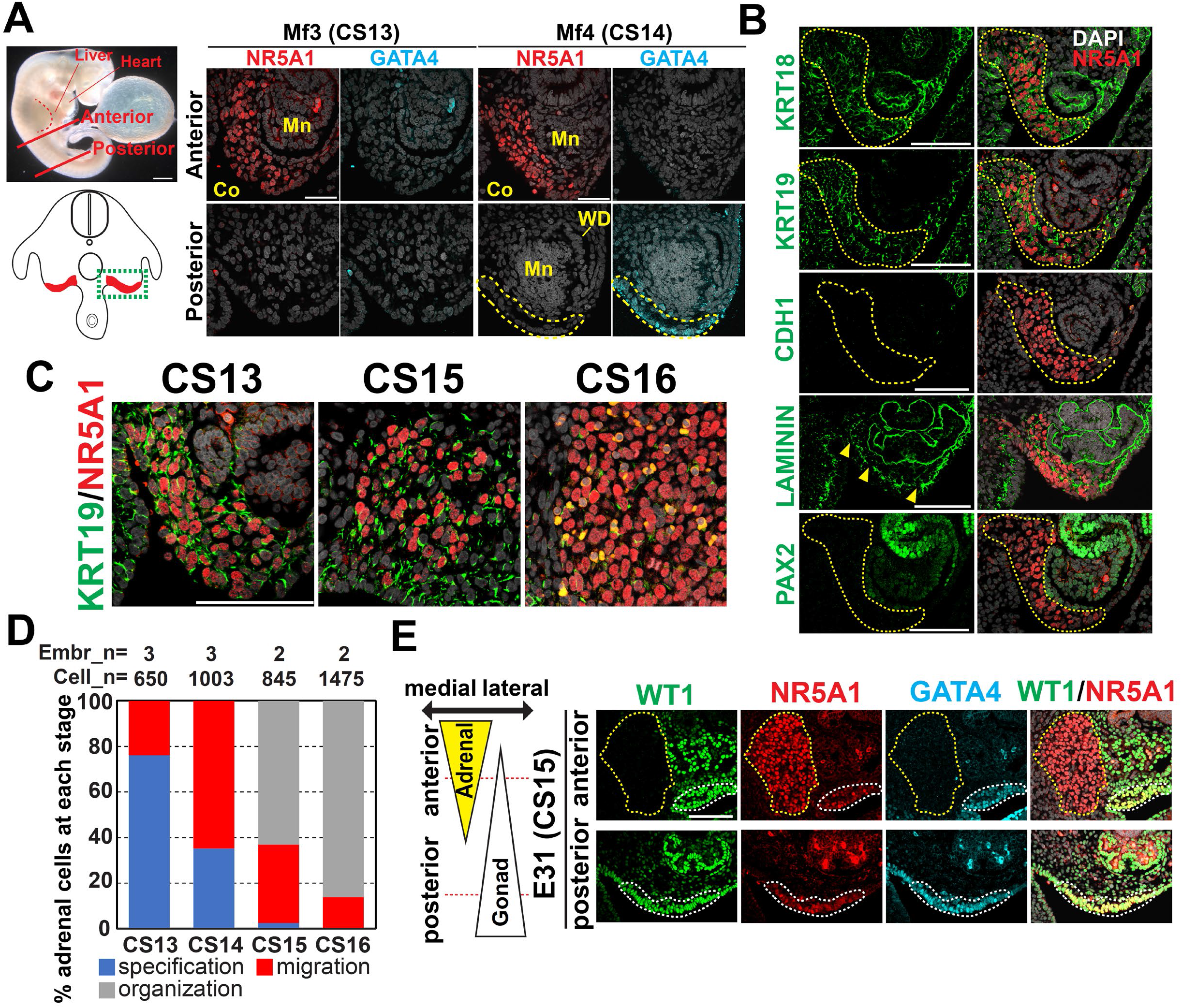
Specification of the adrenal cortex in cynomolgus monkeys. (A) (left top) Brightfield image of a cynomolgus embryo at CS13. The red dotted line denotes the forelimb bud. The red solid lines indicate the approximate planes where the transverse sections were obtained for IF analysis. (left bottom) Schematic illustration of transverse sections. Dotted lines outline the coelomic angle where IF images were taken. (right) IF images of the coelomic angle from cynomolgus embryos at CS13 (left, Mf3) and CS14 (right, Mf4) for NR5A1 (red) and GATA4 (cyan), merged with DAPI (white). Co, coelom; Mn, mesonephros; WD, Wolffian duct. (B) IF images of the coelomic angle (embryos: CS13) for the indicated markers (green) (left). Merged images with NR5A1 (red) and DAPI (white) are shown (right). The yellow dotted lines outline NR5A1^+^ adrenogenic CE. The arrowheads denote regions with basement membrane disruption. Bar, 100 μm. (C) IF images of the adrenocortical cells predominantly in the specification (CS13, left), migration (CS15, middle) or organization stage (CS16, right) for KRT19 (green) and NR5A1 (red), merged with DAPI (white). Bar, 20 μm. (D) Percentage of adrenocortical cells at the indicated morphogenetic stages (specification, migration and organization) for each embryonic stage. Numbers of embryos (Embr_n) and counted cells (cell_n) are shown. (E) (left) Schematic depicting the topological orientation of the adrenal and gonadal linage in a cynomolgus embryo at E31 (CS15). The red dotted lines denote the approximate regions where the sections were made. (right) IF images of the coelomic angles at the representative regions for WT1 (green), NR5A1 (red) and GATA4 (cyan), and merged images for WT1, NR5A1 and DAPI. The yellow and white dotted lines outline the adrenal primordium and gonad progenitor, respectively.

**Fig. S6.**
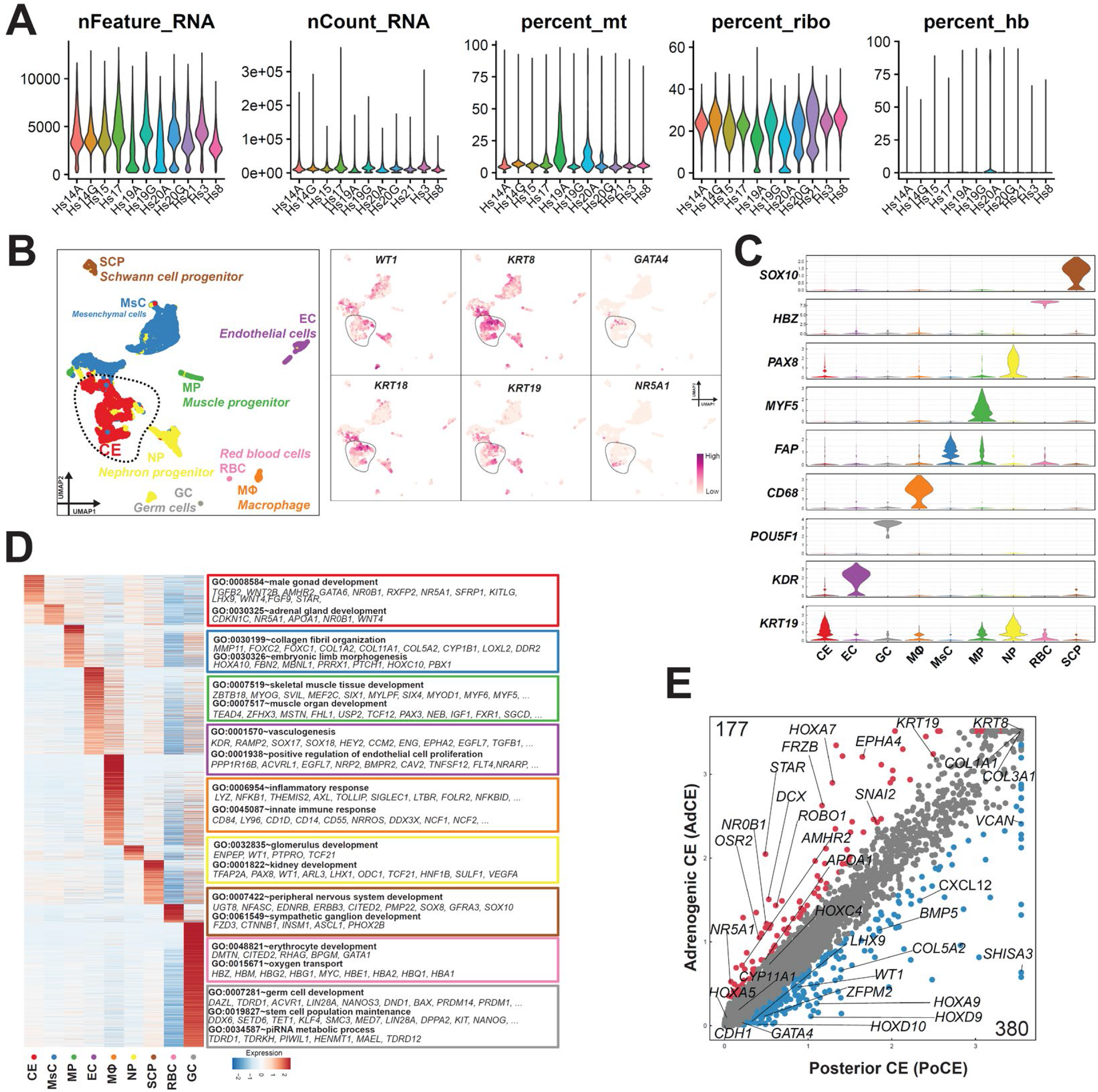
Quality control of scRNA-seq data and analyses of cell types in the human urogenital ridge at 4 wpf. (A) Quality control analyses of all human samples used in this study. (B) (left) UMAP plot showing different cell clusters in a human embryo at 4 wpf. Cell clusters are annotated on the basis of marker genes. A cluster representing the CE is encircled and isolated for sub-clustering in Fig. 3. (right) Expression of key marker genes for CE. EC, endothelial cells; GC, germ cells; MΦ, macrophage; MsC, mesenchymal cells; MP, muscle progenitor; NP, nephron progenitor; RBC, red blood cells; SCP, Schwann cell progenitor. (C) Violin plot showing the expression of key marker genes in the indicated cell types used for annotating cell clusters in (B). (D) (left) Heatmap showing differentially expressed genes (DEGs) among cell clusters. (right) Enriched GO terms and representative genes. Gene expression is Z-score normalized in each row. (E) Scatterplot of pair-wise comparison between adrenogenic CE and posterior CE. DEGs defined as genes with log2 fold change >1, *p*-value <0.01 and FDR <0.01 are shown in colored dots. Key genes and the numbers of DEGs are shown.

**Fig. S7.**
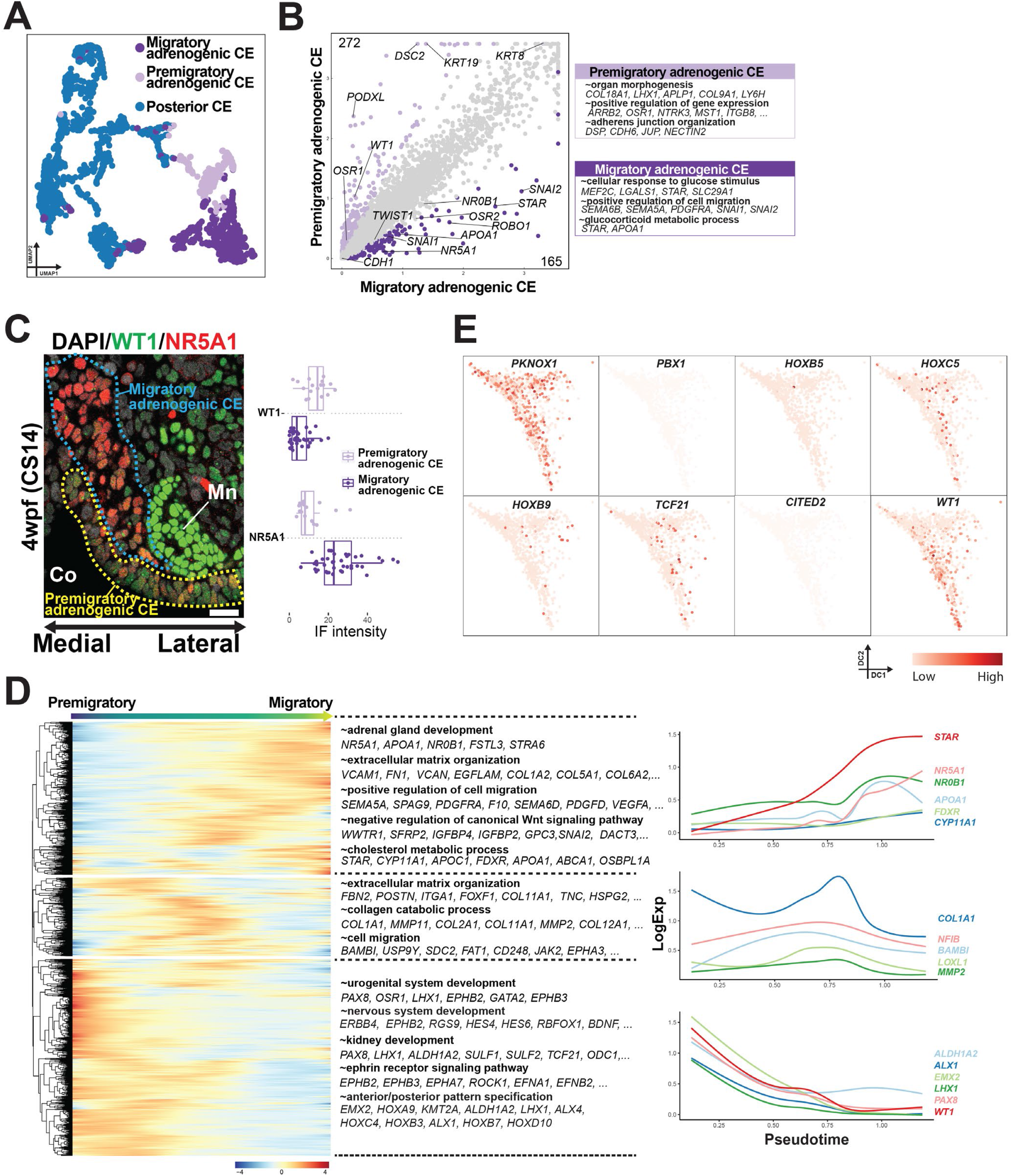
Gene expression dynamics during the migration of adrenogenic coelomic epithelium. (A) Subclustering of the coelomic epithelium (CE) (defined in Fig. S6B) into posterior CE, premigratory adrenogenic CE and migratory adrenogenic CE, projected on a UMAP plot. (B) (left) Scatterplot of pair-wise comparison between premigratory adrenogenic CE and migratory adrenogenic CE. DEGs defined as genes with log2 fold change >1, *p*-value<0.01 and FDR <0.01 are shown in colored dots. Key genes and the numbers of DEGs are shown. (right) List of DEGs and enriched GO terms. (C) (left) Merged IF image for WT1 (green), NR5A1(red) and DAPI (white), showing the histologic distribution of premigratory adrenogenic CE and migratory adrenogenic CE in a transverse section of a 4 wpf human embryo. (right) Quantification of fluorescence intensity, showing that the adrenogenic CE loses WT1 while gaining NR5A1 expression during the transition. (D) (left) Transcriptome dynamics from premigratory adrenogenic CE to migratory adrenogenic CE, corresponding to Fig. 3D. The top 2000 high variable genes are hierarchically clustered with different patterns along pseudotime. Enriched GO terms are listed at right. (right) Expression dynamics of key variable genes in each cluster at left, aligned along pseudotime.

**Fig. S8.**
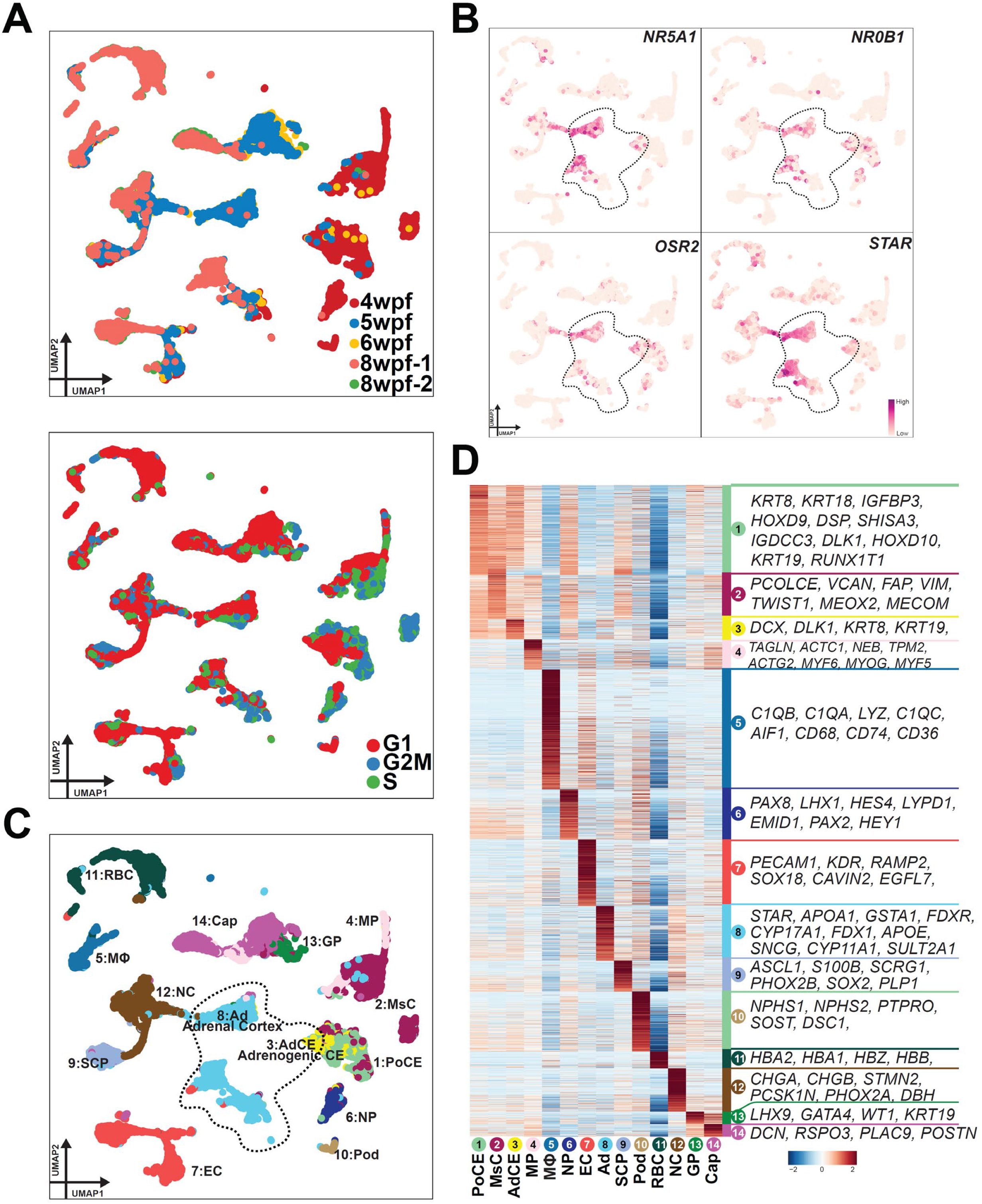
scRNA-seq analyses to define adrenal lineages in human embryos. **(A)** UMAP plot showing all cells from human embryonic adrenal glands (5–8 wpf) and a urogenital ridge (4 wpf) used to isolate the adrenal lineage for the trajectory analysis in Fig. 3F. Each dot represents a single cell, colored according to sample origin (top) and cell cycle scoring (bottom). **(B)** Expression of key marker genes projected on a UMAP plot to define adrenal lineages. Cells outlined by dotted lines are annotated as having adrenal lineage in (C) and were isolated for further sub-clustering in Fig. 3F. **(C)** Cell clusters and their annotations, projected on the UMAP plot used in (A, B). PoCE, posterior CE; MsC, mesenchymal cells; AdCE, adrenogenic CE; MP, muscle progenitor; MΦ, macrophage; NP, nephron progenitor; EC, endothelial cells; Ad, adrenal cortex; SCP, Schwann cell progenitor; Pod, podocytes; RBC, red blood cells; NC, neuroendocrine cells; GP, gonad progenitor; Cap, capsular cells. **(D)** Heatmap of DEGs among the cell clusters. Representative genes are listed at right. Gene expression is Z-score normalized in each row.

**Fig. S9.**
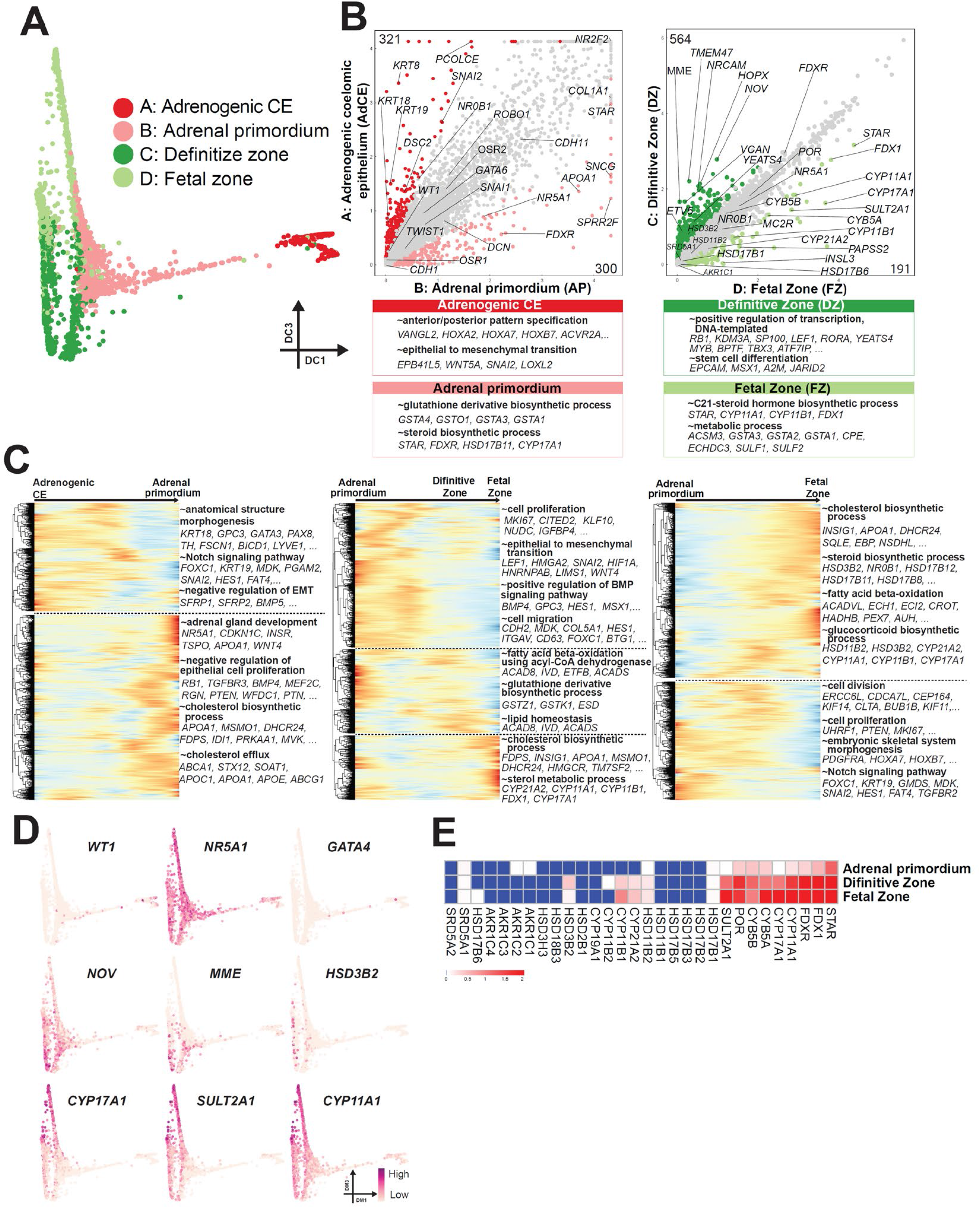
Gene expression dynamics during early adrenal development in humans. (A) 2D diffusion map using the first and third diffusion components and the same clusters as in Fig. 3F. (B) (top) Scatterplots comparing the averaged gene expression levels between adrenogenic CE (cluster A) and adrenal primordium (cluster B) (left), or definitive zone (cluster C) and fetal zone (cluster D) (right). Differentially expressed genes (DEGs) upregulated more than 2-fold (*p*-value <0.01 and FDR <0.01) are shown. Key genes are annotated. (bottom) GO analyses of DEGs. Representative genes in each GO category are shown. (C) Heatmap showing transcriptomic dynamics during early adrenal development. Each row represents a gene, and each column represents a cell, which is ordered by the pseudotime defined in Fig. 3H. Trajectories partitioned in three branches are shown separately (adrenogenic CE to adrenal primordium, left; adrenal primordium to fetal zone via definitive zone, middle; adrenal primordium to fetal zone, right). The top 2000 variable genes are hierarchically clustered. Representative genes and their enriched GO terms for variable genes clustered on the basis of expression patterns are shown at right. (D) Expression of key genes projected on the 2D diffusion map, as in A. (E) Heatmap showing the averaged expression of key genes associated with steroidogenesis in the adrenal primordium, definitive zone and fetal zone. Color indicates log normalized expression.

**Fig. S10.**
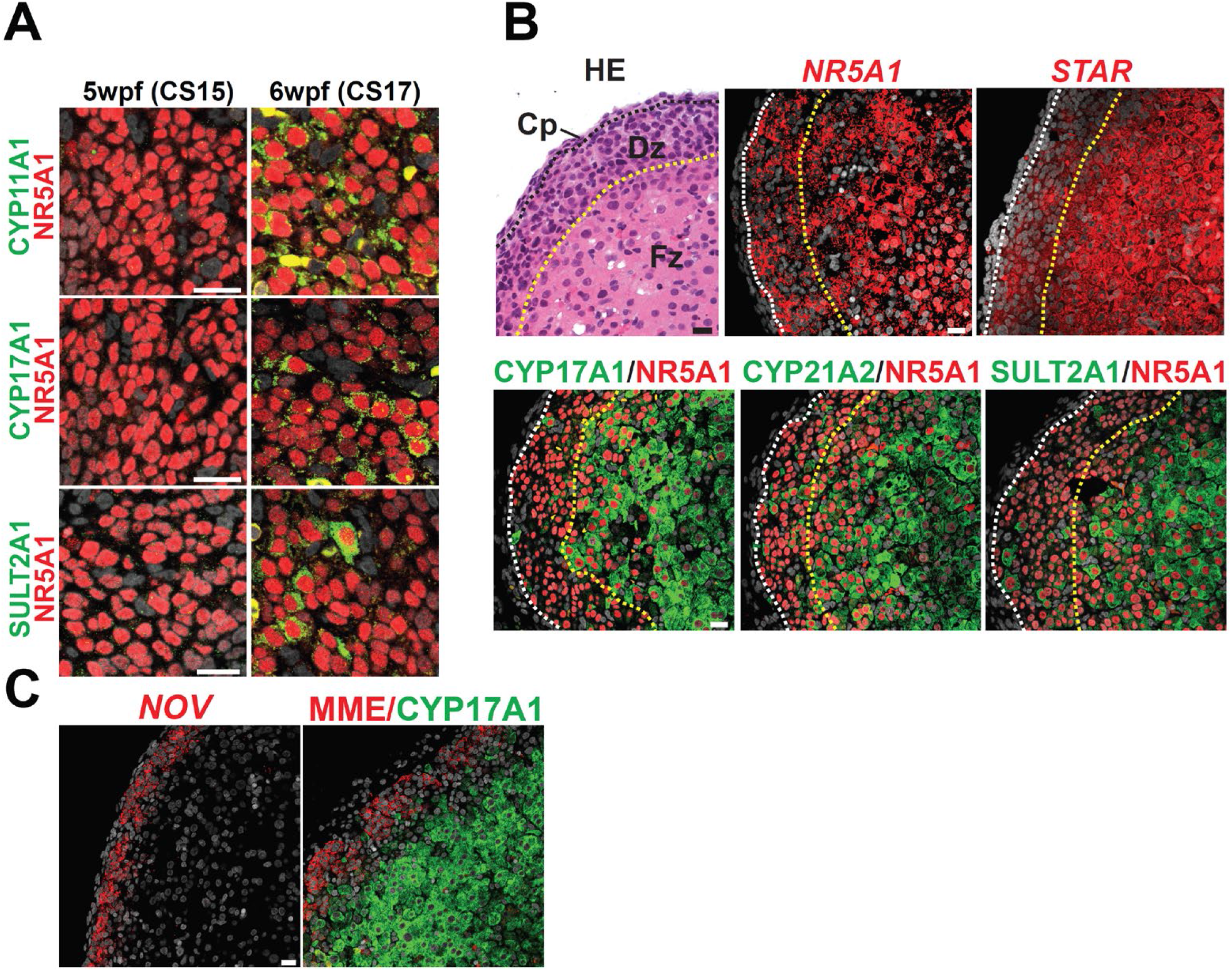
Initiation of steroidogenesis and zonation in developing human adrenal glands. (A) IF of transverse sections of human embryos at the indicated stage for NR5A1 (red) and key steroidogenesis genes (CYP11A1, CYP17A1 and SULT2A) (green) merged with DAPI (white). Bar, 20 μm. (B) (top left) H&E image of a cross section of human fetal adrenals at 8 wpf. (top right) ISH images of the same embryo for *NR5A1* or *STAR* (red), merged with DAPI (white). (bottom) IF images of the same embryo for key steroidogenesis genes (CYP11A1, CYP17A1 and SULT2A) (green) merged with NR5A1 (red) and DAPI (white). The white dotted lines separate the capsule and definitive zone. The yellow dotted lines separate the definitive zone from the fetal zone. Bar, 20 μm. (C) (left) ISH on a section of the adrenals at 8 wpf, showing specific expression of *NOV* (Red) in the definitive zone. (right) IF on the neighboring section for MME (red) and CYP17A1 (green), highlighting the definitive zone and fetal zone, respectively. Merged images with DAPI are shown. Bar, 20 μm.

**Fig. S11.**
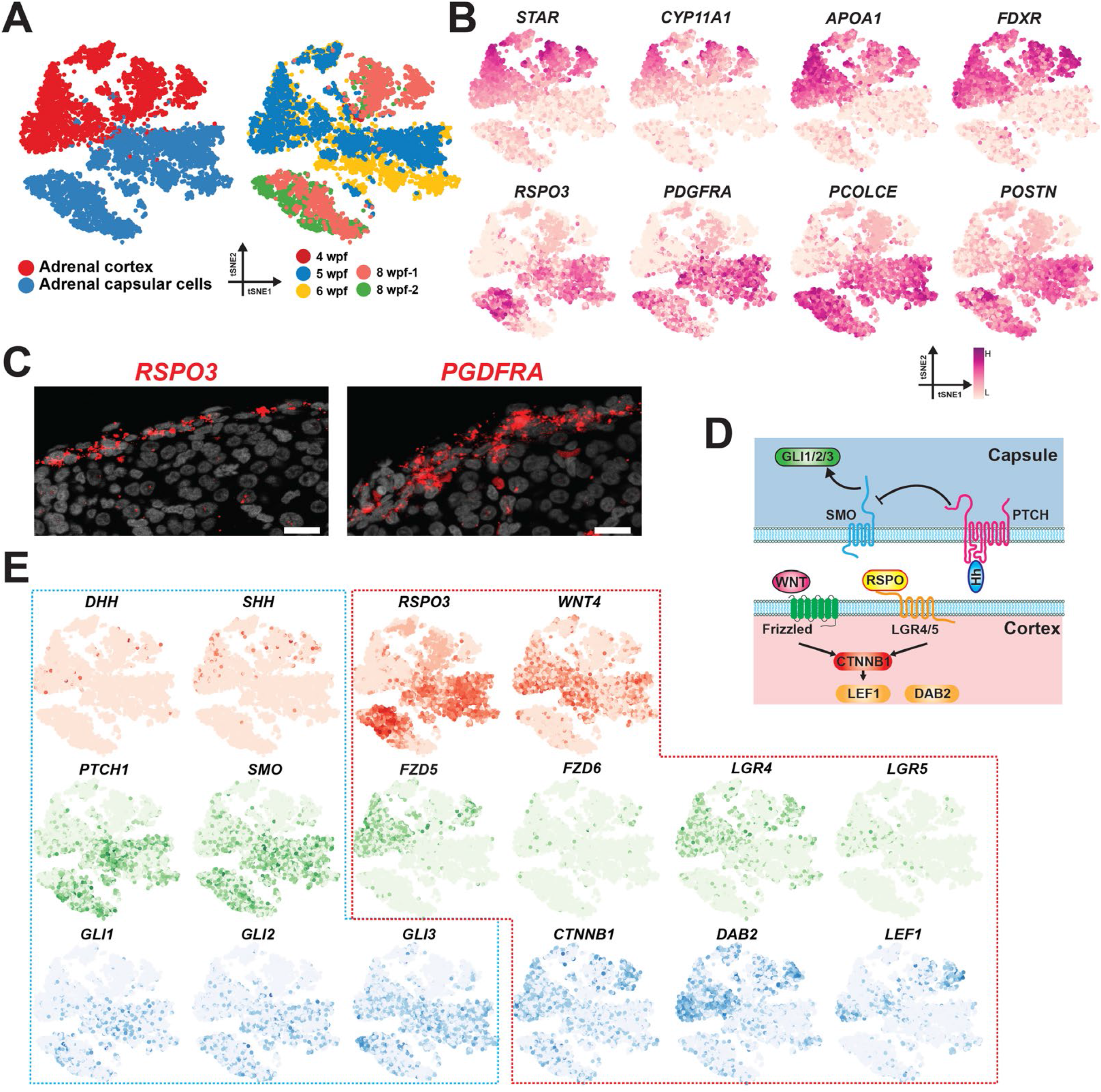
Receptor-ligand interaction between the cortex and capsule in human fetal adrenals. (A) (left) Adrenal cortex (clusters 8 in Fig. S8C) and capsular cells (cluster 14 in Fig. S8C), projected on the *t*-distributed stochastic neighbor embedding (tSNE) plot. (right) Sample origins projected on the same tSNE plot. (B) Expression of key marker genes used for annotation of the adrenal cortex and capsular cells. (C) ISH on the paraffin section of human fetal adrenal cortex (8 wpf) for *RSPO3* and *PDGFRA* (red), merged with DAPI (white). Bar, 20 μm. (D) Diagram depicting the interactions between the cortex and the capsule. (E) Expression of ligands (top), receptors (middle) and target genes (bottom), projected on the tSNE in (A). Hedgehog and WNT pathways are outlined in blue and red dotted lines, respectively.

**Fig. S12.**
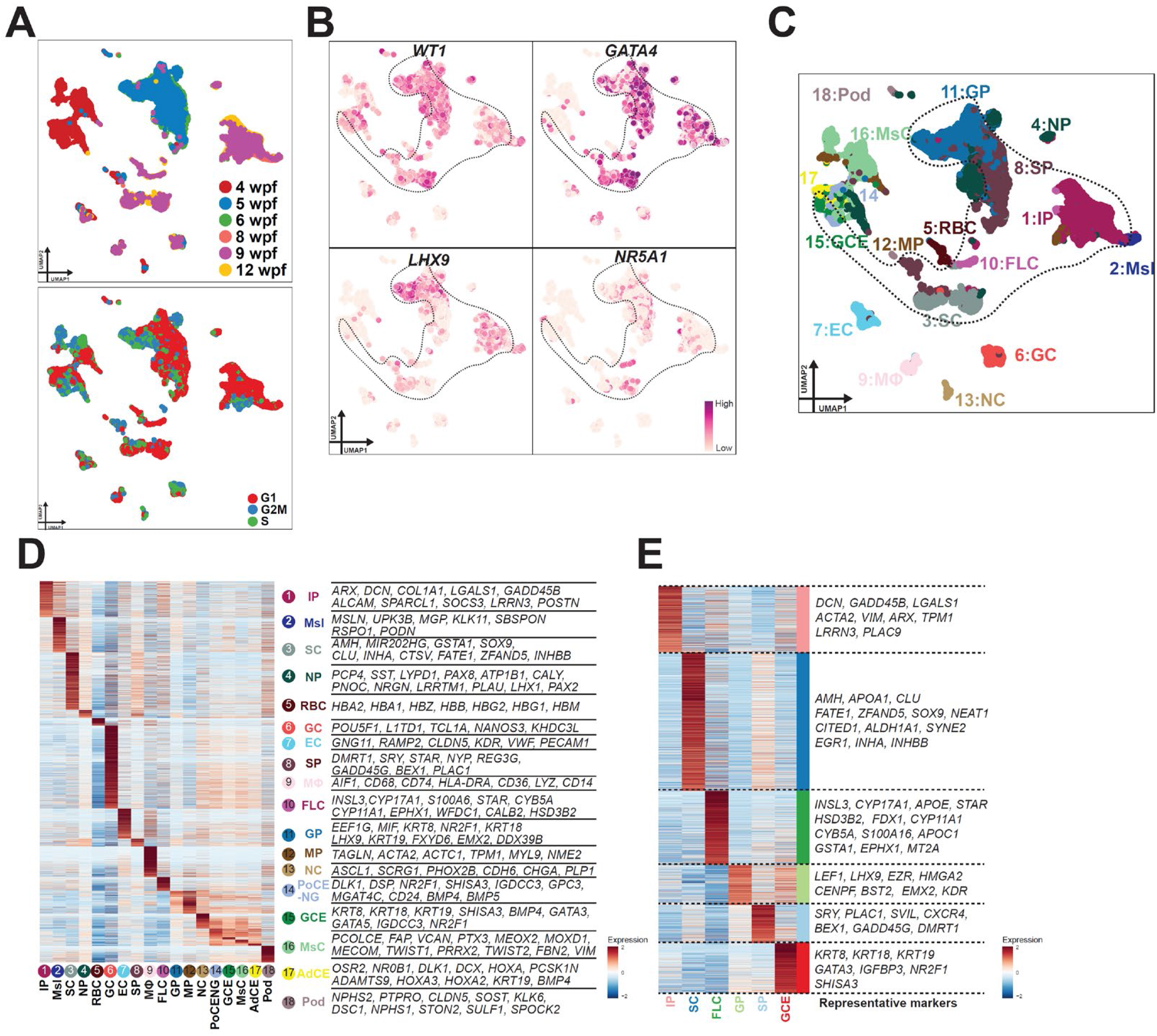
scRNA-seq analyses to define gonadal lineages in human embryos. (A) UMAP plot showing all cells from human embryonic testicular cells (5–12 wpf) and the urogenital ridge (4 wpf) used to isolate the testicular lineage for the trajectory analysis in Fig. 4A. Each dot represents a single cell, colored according to its sample origin (top) and cell cycle scoring (bottom). (B) Expression of key marker genes of the gonadal (testicular) lineage, projected on the UMAP plot in (A). Cells in circles are annotated as having gonadal lineage and isolated for trajectory analysis in Fig. 4A. (C) Cell clusters and their annotations, projected on a UMAP plot. Cells are colored according to cell cluster. IP, interstitial progenitor; Msl, mesothelium; SC, Sertoli cells; NP, nephron progenitor; RBC, red blood cells; GC, germ cells; EC, endothelial cells; SP, Sertoli progenitor; MΦ, macrophage; FLC, fetal Leydig cells; GP, gonad progenitor; MP, muscle progenitor; NC, neuroendocrine cells; GCE, gonadogenic CE; MsC, mesenchymal cells; Pod, podocytes. (D) Heatmap showing the averaged expression pattern of DEGs among the cell clusters (left) and key genes (right). Gene expression is Z-score normalized in each row. PoCE- NG, non-gonadal posterior CE; AdCE, adrenogenic CE. (E) Heatmap showing the averaged expression pattern of DEGs among different cell clusters of gonadal lineages identified in Fig. 4A (left) and representative genes (right). Gene expression is Z-score normalized by row.

**Fig. S13.**
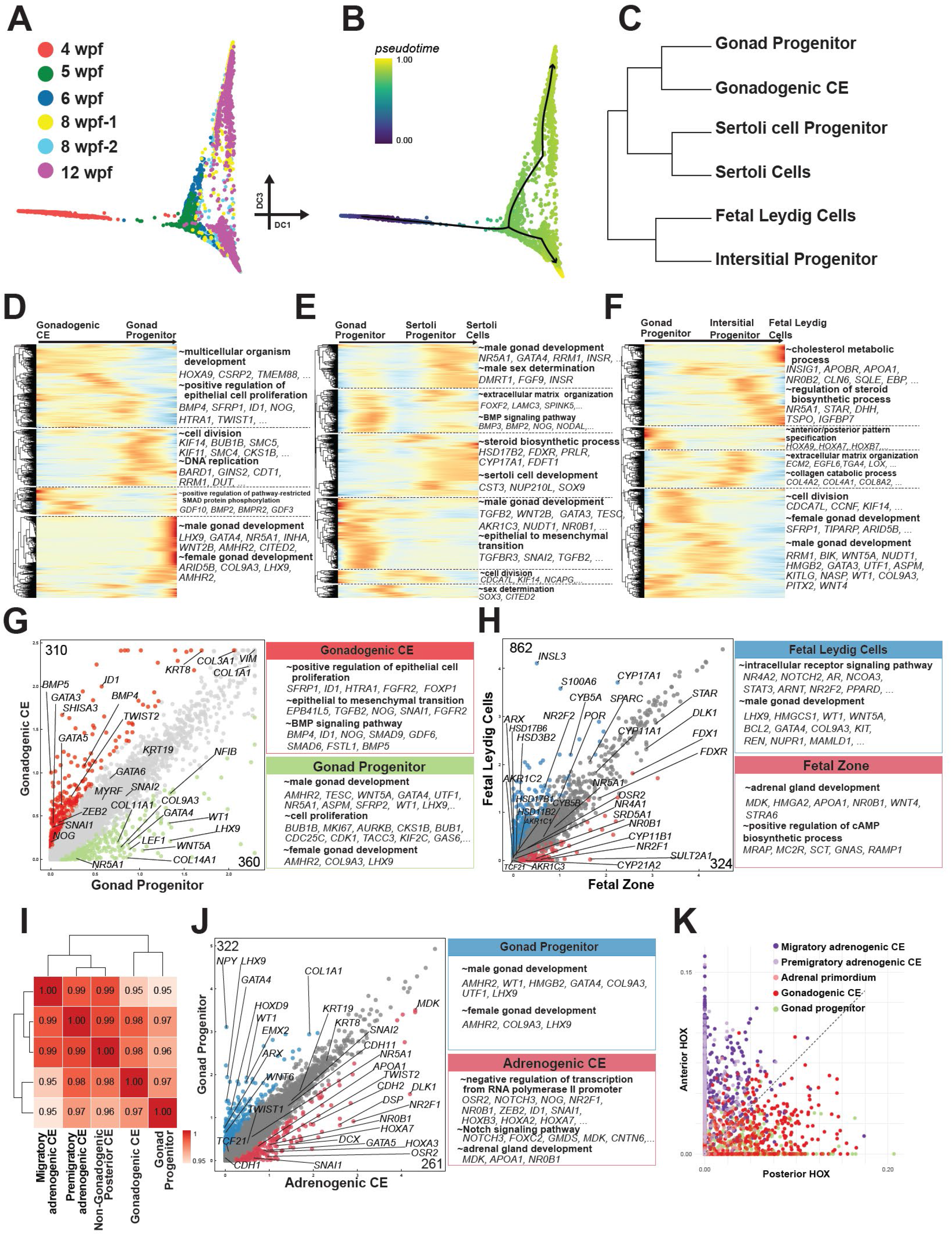
Characterization of developing gonadal lineages and comparison with adrenal lineages in humans. (A) Projection of sample origins on the diffusion map used in Fig. 4A. (B) Projection of pseudotime and fitted principal curves on the diffusion map. (C) Hierarchical clustering dendrogram showing the relationships among the cell clusters. The correlation matrix was computed with the Spearman method. (D, E, F) Heatmap showing gene expression dynamics along lineage trajectories (defined in B) from gonadogenic CE to gonad progenitor (D), from gonad progenitor to Sertoli cells (E), and from gonad progenitor to fetal Leydig cells (F). Each row is a gene, each column is a cell, and cells are ordered by pseudotime as in (B). The top 2000 high variable genes are hierarchically clustered. Gene expression is Z-score normalized by row. (G) Scatterplot comparing the averaged gene expression levels between gonadogenic CE and gonad progenitor. DEGs upregulated in gonadogenic CE (red) and gonad progenitor (green) (2-fold change, *p*-value <0.01 and FDR <0.01) are highlighted, and key genes are annotated. (right) GO analyses of the DEGs. Representative genes in each GO category are shown. (H) (left) Scatterplot comparison of the averaged values of DEGs between fetal Leydig cells and the fetal zone. Genes upregulated by more than 2-fold (*p*-value <0.01, FDR <0.01) in the indicated cell clusters are plotted with colors. Key genes and the numbers of DEGs are shown. (right) Over-represented GO terms and representative genes in each GO category are shown. (I) Spearman correlation and hierarchical clustering of the cell clusters, showing that the gonad progenitor is close to gonadogenic CE rather than adrenogenic CE. (J) (left) Scatterplot comparison of the averaged values of DEGs between gonad progenitor and adrenogenic CE. Genes upregulated by more than 2-fold (*p*-value <0.01, FDR <0.01) in the indicated cell clusters are plotted with colors. Key genes and the numbers of DEGs are shown. (right) Over-represented GO terms and representative genes in each GO category are shown. (K) Area under the curve (AUC) scoring of *HOX* genes in the indicated cell types. Each dot represents a single cell, colored according to the cell clusters annotated in Fig. 3D, F and 4A. Cells in the gonadal lineage (gonadogenic CE, gonad progenitor) are biased toward higher posterior HOX scores, whereas cells in the adrenal lineage (migratory adrenogenic CE, premigratory adrenogenic CE, adrenal primordium) are biased toward higher anterior HOX scores.

### Supplementary Tables (Separate files)

Table S1. Human and cynomolgus embryos used in this study.

Table S2. Differentially expressed genes across cell clusters in the human urogenital ridge at 4 wpf.

Table S3. Differentially expressed genes between the pre-migratory and migratory adrenogenic coelomic epithelium.

Table S4. Differentially expressed genes across cell clusters present in human adrenal glands (5-12 wpf) and the urogenital ridge (4 wpf).

Table S5. Differentially expressed genes across cell clusters in human adrenal lineages.

Table S6. Differentially expressed genes across cell clusters in human testes and the urogenital ridge (4 wpf).

## REFERENCES

1. E. Pignatti, C. E. Flück, Adrenal cortex development and related disorders leading to adrenal insufficiency. Molecular and Cellular Endocrinology. 527, 111206 (2021).

2. H. Ishimoto, R. B. Jaffe, Development and Function of the Human Fetal Adrenal Cortex: A Key Component in the Feto-Placental Unit. Endocrine Reviews. 32, 317– 355 (2011).

3. H. Fabbri-Scallet, L. M. de Sousa, A. T. Maciel-Guerra, G. Guerra-Júnior, M. P. de Mello, Mutation update for the NR5A1 gene involved in DSD and infertility. Human Mutation. 41, 58–68 (2020).

4. M. Zubair, S. Ishihara, S. Oka, K. Okumura, K. Morohashi, Two-Step Regulation of Ad4BP/SF-1 Gene Transcription during Fetal Adrenal Development: Initiation by a Hox-Pbx1-Prep1 Complex and Maintenance via Autoregulation by Ad4BP/SF-1 . Molecular and Cellular Biology. 26, 4111–4121 (2006).

5. M. Zubair, K. L. Parker, K. Morohashi, Developmental Links between the Fetal and Adult Zones of the Adrenal Cortex Revealed by Lineage Tracing. Molecular and Cellular Biology. 28, 7030–7040 (2008).

6. R. Bandiera, V. P. I. Vidal, F. J. Motamedi, M. Clarkson, I. Sahut-Barnola, A. 16 vonGise, W. T. Pu, P. Hohenstein, A. Martinez, A. Schedl, WT1 Maintains Adrenal- Gonadal Primordium Identity and Marks a Population of AGP-like Progenitors within the Adrenal Gland. Developmental Cell. 27, 5–18 (2013).

7. O. Hatano, A. Takakusu, M. Nomura, K. I. Morohashi, Identical origin of adrenal cortex and gonad revealed by expression profiles of Ad4BP/SF-1. Genes to Cells. 1, 663–671 (1996).

8. K. Sasaki, A. Oguchi, K. Cheng, Y. Murakawa, I. Okamoto, H. Ohta, Y. Yabuta, C. Iwatani, H. Tsuchiya, T. Yamamoto, Y. Seita, M. Saitou, The embryonic ontogeny of the gonadal somatic cells in mice and monkeys. Cell Reports. 35, 109075 (2021).

9. P. Val, J.-P. Martinez-Barbera, A. Swain, Adrenal development is initiated by Cited2 and Wt1 through modulation of Sf-1 dosage. Development. 134, 2349–2358 (2007).

10. N. A. Hanley, S. G. Ball, M. Clement-Jones, D. M. Hagan, T. Strachan, S. Lindsay, S. Robson, H. Ostrer, K. L. Parker, D. I. Wilson, Expression of steroidogenic factor 1 and Wilms’ tumour 1 during early human gonadal development and sex determination. Mechanisms of Development. 87, 175–180 (1999).

11. N. A. Hanley, W. E. Rainey, D. I. Wilson, S. G. Ball, K. L. Parker, Expression Profiles of SF-1, DAX1, and CYP17 in the Human Fetal Adrenal Gland: Potential Interactions in Gene Regulation. Molecular Endocrinology. 15, 57–68 (2001).

12. M. E. Sucheston, M. S. Cannon, Development of zonular patterns in the human adrenal gland. Journal of Morphology. 126, 477–491 (1968).

13. J. Jirasek, Human Fetal Endocrines (Martinus Nijhoff, London, United Kingdom, 1980).

14. R. Lyraki, A. Schedl, Adrenal cortex renewal in health and disease. Nature Reviews Endocrinology 2021 17:7. 17, 421–434 (2021).

15. H. Ishimoto, R. B. Jaffe, Development and Function of the Human Fetal Adrenal Cortex: A Key Component in the Feto-Placental Unit (2011), doi:10.1210/er.2010-0001.

16. Y.-C. Hu, L. M. Okumura, D. C. Page, Gata4 Is Required for Formation of the Genital Ridge in Mice. PLoS Genetics. 9, e1003629 (2013).

17. G. R. Dressler, A. S. Woolf, Pax2 in development and renal disease. International Journal of Developmental Biology. 43, 463–468 (2003).

18. N. A. Hanley, S. G. Ball, M. Clement-Jones, D. M. Hagan, T. Strachan, S. Lindsay, S. Robson, H. Ostrer, K. L. Parker, D. I. Wilson, Expression of steroidogenic factor 1 and Wilms’ tumour 1 during early human gonadal development and sex determination. Mechanisms of Development. 87, 175–180 (1999).

19. J. Yang, P. Antin, G. Berx, C. Blanpain, T. Brabletz, M. Bronner, K. Campbell, A. Cano, J. Casanova, G. Christofori, S. Dedhar, R. Derynck, H. L. Ford, J. Fuxe, A. García de Herreros, G. J. Goodall, A. K. Hadjantonakis, R. J. Y. Huang, C. Kalcheim, R. Kalluri, Y. Kang, Y. Khew-Goodall, H. Levine, J. Liu, G. D. Longmore, S. A. Mani, J. Massagué, R. Mayor, D. McClay, K. E. Mostov, D. F. Newgreen, M. A. Nieto, A. Puisieux, R. Runyan, P. Savagner, B. Stanger, M. P. Stemmler, Y. Takahashi, M. Takeichi, E. Theveneau, J. P. Thiery, E. W. Thompson, R. A. Weinberg, E. D. Williams, J. Xing, B. P. Zhou, G. Sheng, Guidelines and definitions for research on epithelial–mesenchymal transition. Nature Reviews Molecular Cell Biology. 21, 341–352 (2020).

20. J. Guo, E. Sosa, T. Chitiashvili, X. Nie, E. J. Rojas, E. Oliver, K. Plath, J. M. Hotaling, J. B. Stukenborg, A. T. Clark, B. R. Cairns, Single-cell analysis of the developing human testis reveals somatic niche cell specification and fetal germline stem cell establishment. Cell Stem Cell. 28, 764–778.e4 (2021).

21. Crowder, RE., The development of the adrenal gland in man, with special reference to origin and ultimate location of cell types and evidence in favor of the “cell migration” theory. Contemp Embryol. 251, 195–209 (1957).

22. U. U. Uotila, The early embryological development of the fetal and permanent adrenal cortex in man. The Anatomical Record. 76, 183–203 (1940).

23. J. Deschamps, D. Duboule, Embryonic timing, axial stem cells, chromatin dynamics, and the Hox clock. Genes and Development. 31 (2017), pp. 1406–1416.

24. R. Neijts, S. Simmini, F. Giuliani, C. van Rooijen, J. Deschamps, Region-specific regulation of posterior axial elongation during vertebrate embryogenesis. Developmental Dynamics. 243, 88–98 (2014).

25. A. Taguchi, R. Nishinakamura, Higher-Order Kidney Organogenesis from Pluripotent Stem Cells. Cell Stem Cell. 21, 730–746.e6 (2017).

26. A. Taguchi, Y. Kaku, T. Ohmori, S. Sharmin, M. Ogawa, H. Sasaki, R. Nishinakamura, Redefining the in vivo origin of metanephric nephron progenitors enables generation of complex kidney structures from pluripotent stem cells. Cell Stem Cell. 14, 53–67 (2014).

27. K. Sasaki, T. Nakamura, I. Okamoto, Y. Yabuta, C. Iwatani, H. Tsuchiya, Y. Seita, R. Nakamura, N. Shiraki, T. Takakuwa, T. Yamamoto, M. Saitou, The Germ Cell Fate of Cynomolgus Monkeys Is Specified in the Nascent Amnion. Developmental Cell. 39, 169–185 (2016).

28. J. Yamasaki, C. Iwatani, H. Tsuchiya, J. Okahara, T. Sankai, R. Torii, Vitrification and transfer of cynomolgus monkey (Macaca fascicularis) embryos fertilized by intracytoplasmic sperm injection. Theriogenology. 76, 33–38 (2011).

29. R. O’Rahilly, F. Muller, Developmental stages in human embryos. Carnegie Inst Washington Publ. 637 (1987).

